# Allele surfing causes maladaptation in a Pacific salmon of conservation concern

**DOI:** 10.1101/2022.11.10.515805

**Authors:** Quentin Rougemont, Thibault Leroy, Eric B. Rondeau, Ben Koop, Louis Bernatchez

## Abstract

How various factors, including demography, recombination or genome duplication, may impact the efficacy of natural selection and the burden of deleterious mutations,is a central question in evolutionary biology and genetics. In this study, we show that key evolutionary processes, including variations in *i*) effective population size (*N_e_*) *ii*) recombination rates and *iii*) chromosome inheritance, have influenced the genetic load and efficacy of selection in Coho salmon (*Oncorhynchus kisutch*), a widely distributed salmonid species on the west coast of North America. Using whole genome resequencing data from 14 populations at different migratory distances from their southern glacial refugium, we found evidence supporting gene surfing, wherein reduced *N_e_* at the postglacial recolonization front, leads to a decrease in the efficacy of selection and a surf of deleterious alleles in the northernmost populations. Furthermore, our results indicate that recombination rates play a prime role in shaping the load along the genome. Additionally, we identified variation in polyploidy as a contributing factor to within-genome variation of the load. Overall, our results align remarkably well with expectations under the nearly neutral theory of molecular evolution. We discuss the fundamental and applied implications of these findings for evolutionary and conservation genomics.

**Author Summary:** Understanding how historical processes, such as past glaciations, may have impacted variations in population size and genetic diversity along the genome is a fundamental question in evolution. In this study, we investigated how recent postglacial demographic expansion has affected the distribution of deleterious genetic variants and the resulting deleterious mutation load in Coho salmon (*Oncorhynchus kisutch*), throughout its native range in North America. By sequencing the entire genome of 71 Coho salmon, we reveal that postglacial expansion has led to allele surfing, a process where alleles increase in frequency in populations that are expanding or colonizing new environments. Here, allele surfing resulted in an increased deleterious mutation load at the colonization front. Furthermore, we demonstrated that the efficacy of natural selection scales with variation in effective population size among populations. We showed that the specific genomic features of Coho salmon, namely variation in local recombination rate and variation in chromosomal inheritance, strongly impacted the segregation of deleterious mutations.

## Introduction

Demographic events, including population contractions, expansions and secondary contacts have profound impacts on the spatial distribution of genetic diversity within species [1]. Range expansions can be accompanied by multiple founder events when few individuals colonize new areas. At the wave front, this process results in increased genetic drift and a reduced effective population size [2,3]. This has multiple consequences genome-wide including (i) extended linkage disequilibrium [4], (ii) a loss of genetic diversity and (iii) increased levels of genetic differentiation [2,3,5]. At the wavefront, increased genetic drift favors allele surfing, a process that will increase the relative proportion of neutral, deleterious or advantageous mutations that will fix in the population [5–7]. Such allele surfing is expected to have two main negative consequences at the wave front: *i*) an increase of the deleterious load (called expansion load [8,9]) and *ii*) a loss of adaptive potential [10].

First, allele surfing through increased genetic drift can overwhelm the effect of weak purifying selection, resulting in an increase of the deleterious load [11]. Due to the lower efficacy of natural selection in low *N_e_* populations, a greater fraction of slightly deleterious mutations is expected to be present in populations at the wavefront (*i.e.,* elevated ratios of non-synonymous to synonymous nucleotide diversity (π_N_/π_S_) [8,9]). Moreover, under the nearly neutral theory of molecular evolution, the reduced efficacy of purifying selection in small populations should increase the fixation rate of slightly deleterious mutations [12]. On the other hand, given that most deleterious mutations are expected to be recessive, increased homozygosity of deleterious mutations should enable their removal more efficiently through genetic purging, leading to a reduction of the recessive and additive load [13]. To take this into account, a possible strategy is to investigate both the additive (i*.e,.* total) and the recessive (*i.e.,* fixed) deleterious mutation load. [8]. Evidence for an increased recessive genetic load has been reported in humans which has been associated with the Out-of-Africa bottleneck [14,15]. Evidence has also been gathered in expanding populations in plants [16–20]. For instance in *Arabidopsis lyrata,* range expansion has been associated with an increased mutational load [16] and increased linkage disequilibrium [21]. Similar results were observed in *Escherichia coli*[22]. In contrast, a recent study in *A. lyrata* found no evidence of increased load following range expansion [23].

Second, allele surfing is expected to lead to a loss of adaptive potential due to a reduction in the rate of adaptive substitutions given that the supply of new mutations is proportional to *N_e_* [24]. Unless compensatory mutations (*i.e.,* favourable mutations whose probability of occurrence increases to compensate for the fixation of slightly detrimental mutations – themselves with a higher probability of fixation in small populations–), counteract this effect in populations with a low *N_e_* [25], one expects a lower proportion of substitutions driven by positive selection [26]. Support for a decreased rate of adaptation associated with range expansion was documented, particularly in plants [17,20]. For instance, such a pattern was reported in the North American populations of *A. lyrata*[17] [27], but not in the European ones despite a slight increase in their genomic burden [23]. Similarly an increased load is observed in human populations, a result that matches theoretical expectations under the out-of-Africa scenario [28] but with limited evidence of a decrease in the efficacy of natural selection [29]. Beyond application to plants, humans and bacteria, there is a lack of empirical studies documenting the consequences of range expansion on the deleterious mutation load and the efficacy of natural selection.

Geographic variation in the overall mutation load is not the only interesting pattern to investigate; the genomic scale is also important. Both deleterious load and efficacy of selection are expected to vary along the genome depending on the local recombination rate [30,31]. Non-recombining regions are expected to more freely accumulate deleterious mutations, a process called Muller’s Ratchet [32]. Moreover, Hill-Robertson interference, a process whereby competing alleles interfere with each other to become fixed, increases the fixation rate of deleterious variants linked to positively selected sites. Such a process will be exacerbated in the absence of recombination [30,33].

In species that have undergone a whole-genome duplication (WGD) with associated residual polyploidy, variation in chromosomal inheritance and genomic architecture may affect the distribution of deleterious mutations along the genome [34]. By doubling the chromosome number, WGD has major consequences. In the short term, WGD alters gene expression and favours polyploid dysfunction (reviewed in [35]). In the middle to long-term, the progressive return to a diploid state will result in regions of residual tetraploidy. Such regions should display an increased effective population size as compared to the rest of the genome (4 *N_e_*instead of 2 *N_e_*) and therefore a higher efficacy of selection and a reduced genetic load. Similarly, the population scale recombination rate, which also depends on *N_e_*, should favour a higher efficacy of selection and a reduced genetic load. We thus predict that these regions should display a lower genetic load compared to diploid regions of the genome. Finally, theoretical work has shown that dominance of mutation, especially in the recessive case, is an important factor influencing the detection of hard sweeps in polyploid genomes [36]. Dominance may therefore also influence the segregation of deleterious variants in regions of residual tetraploidy. WGD has been associated with a reduced efficacy of purifying selection which may favour the accumulation of transposable elements. Such WGD may in turn have contrasted effects, as observed in plants [37,38]. For instance, a LTR insertion favouring early flowering time (an adaptive trait in harsh environments), was shown to be present in tetraploid populations, but not in diploid populations of *A. arenosa*. In contrast, WGS was shown to be associated with small but significant differences in the load and distribution of fitness effect in the same species.

The Coho salmon (*Oncorhynchus kisutch*) is a Pacific salmon species of major cultural and economical importance, which has severely declined over the last decades (reviewed in [39]). It has undergone further recent declines in multiple parts of its native range [40]. In addition to human-induced perturbations, the species has undergone a series of postglacial expansions from its main glacial refugium (Cascadia/California) with multiple founder events along its route toward Alaska, leading to a pattern of isolation by distance, a gradient of population structure and a decrease in genetic diversity [41–43]. Accordingly, these range expansions may result in an increased expansion load, questioning whether such load may have consequences on the fitness of populations at the expansion front. The Coho salmon offers a unique opportunity to test the role of demography and recombination on the deleterious load and on the efficacy of selection. The Coho salmon belongs to the Salmonidae family, a particularly relevant group of species to investigate the effect of WGD. The common ancestor of all present-day salmonids has undergone a whole-genome duplication event ∼80-100MyA [44]. The genome of salmonid species is still on its path to rediploidization. For instance, approximately 8% of the Coho salmon genome displays residual tetraploidy [41]. Residual tetraploidy offers a unique opportunity to investigate the direct effect of the genomic variation of *N_e_* on the efficacy of selection.

To address these questions, we aimed at testing the following general ypotheses: i) whether demographic expansion and bottlenecks have led to an increased load and decreased efficacy of selection, ii) whether heterogeneous recombination levels shape the within-genome variation of the load as a result of Hill-Robertson effects and iii) whether residual tetraploidy results in an increased efficacy of selection and reduced deleterious load through increased recombination and *N_e_*.

## Results and Discussion

### Strong postglacial population expansion revealed through whole genome sequences

We generated 30X coverage whole genome resequencing data for 71 Coho salmon representing 14 populations distributed from California to some of the most upstream populations of the Porcupine River in Yukon, Canada (**Fig 1A**). We also included several outgroup species namely the Sockeye salmon (*O. nerka*), Chinook salmon (*O. tshawytscha*), Pink salmon *(O. Gorbuscha),* Rainbow trout (*O. mykiss*) and Atlantic salmon (*Salmo salar*). These genomes were used to identify the ancestral and derived alleles in Coho and/or testing correlation(s) between genomic load estimates and recombination rate (**Table S1-S3**).

**Figure 1:**
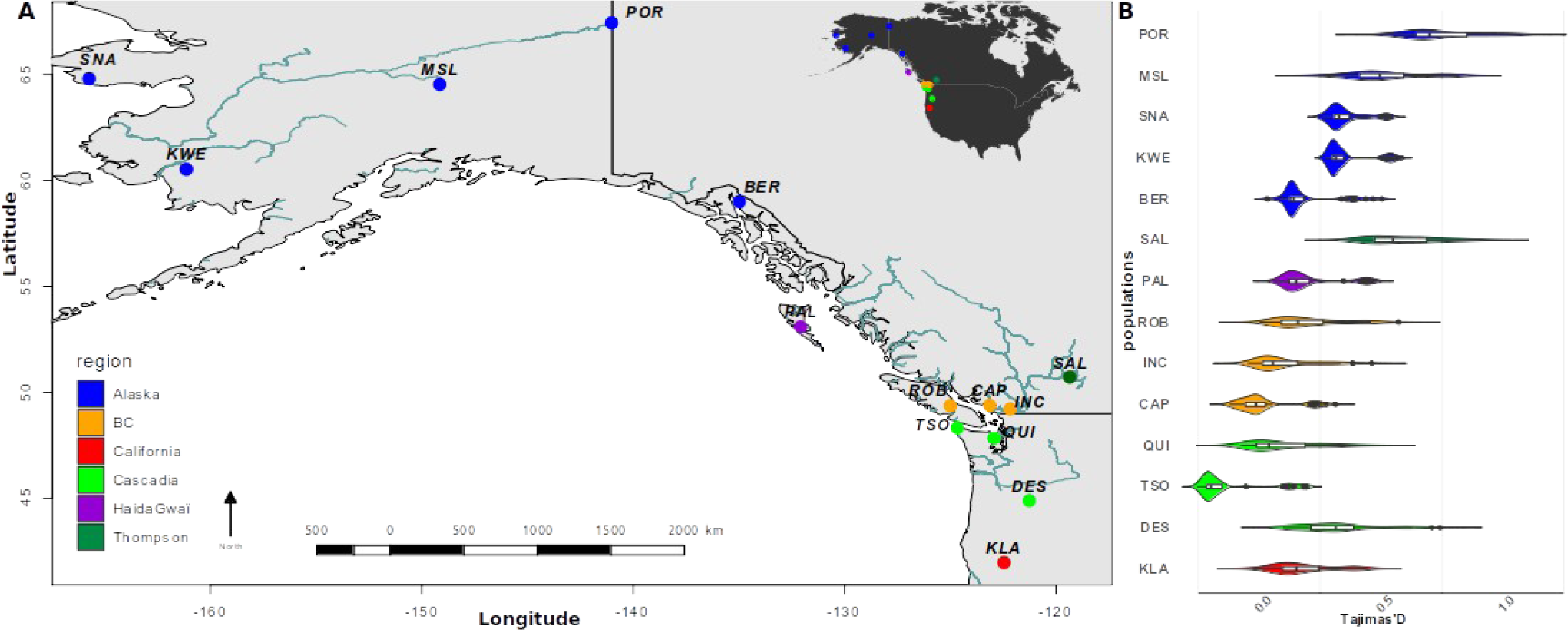
Sampling design and population size change inferred through whole genome sequences. A) Sample site along the Pacific coast of North America. Map produced in R using the World data from Maps package with public domain data available at [47]. B) Average Tajima’s D values computed along the genome and summarized for each sample locality. Populations are ordered from distance to the river mouth and from the South to the North (i) California, (ii) Cascadia, (iii) British-Columbia (BC), (iv) Haida-Gwaii, (v) Salmon River (Fraser Basin) and (vi) Alaska. Abbreviation provided in Table S2). Tajima’s D values vary positively as a function of the distance from the southernmost site (R^2^ = 0.28, p <2e^−16^) and as a function of the distance to the river mouth (R ^2^ = 0.47, p <2e^−16^) reflecting the history of founder events from downstream to upstream sites.

To explore the amount of clustering among individuals a PCA was performed (**Fig S1A, B**). The PCA illustrates that populations are spatially structured following both latitudinal and longitudinal gradients. The first axis strongly correlates with both latitude (Pearson r = 0.97, p < 2e^−16^) and longitude (Pearson r = −0.82, p < 2e^−16^), as expected when space drives population structure [45]. Moreover, admixture analyses (**Fig S1B**) revealed that Coho populations were generally structured at the river/watershed level. These observations corroborate our previous inferences based on lower resolution genomic dataset [42,43].

Under a single refugial expansion scenario, we expect a linear decrease in genetic diversity as a function of the distance from the source. We plotted the distribution of observed heterozygosity (H_o_) as a function of the distance to the southernmost sample and observed a negative correlation with the distance to the southernmost site (R^2^ = 0.77, p<2e^−16^, **Fig S2B**). Next, we used the β_ST_ coefficient to identify the most likely ancestral populations from which the expansion could have started. β_ST_ can account for the non-independence among populations and negative values are indicative of ancestral populations [46]. This metric was also positively correlated with distance to the southernmost site (R^2^ = 0.75, p < 2e^−16^, **Fig S2A**).

The Salmon River population from the Thompson area (dark green point, **Fig 1A** and **Fig S2A**) exhibits lower diversity levels, as compared to all non-Alaskan populations. Excluding this outlier river, revealed even stronger correlations, with a correlation with β_ST_ and H_O_ respectively raising to R^2^ = 0.88 and R^2^ = 0.90 (p <2e^−16^ for both tests), suggesting an overall clear spatial pattern across the populations sampled, with the notable exception of this specific population which had a different demographic trajectory in this population, which seems consistent with a strong bottleneck and inbreeding [41].

Another indicator of changes in population sizes is an increase in Tajima’s D values at the genome-wide level. We tested the hypothesis of increased Tajima’s D values as a function of (i) the traveled distance from the southernmost site (reflecting northward postglacial expansion) and (ii) the distance to the sea only (reflecting upstream directed founder events). Results are consistent with a signal of population expansion along the colonization axis from the south to the north based on Tajima’s D values (R ^2^ =0.28, p< 2e^−16^, **Fig 1B**). Distance to the sea is also highly significant (R^2^ *=* 0.47, p <2e^−16^ for Tajima’s D). This suggests that more genetic drift occurs in upstream populations e.g. Porcupine (POR), Mile Slough (MSL), Thompson (SAL) and Deschutes (DES).

Another hallmark of population bottlenecks and genetic drift associated with founder events is an increase in linkage disequilibrium across the genome [4]. We tested this hypothesis by measuring linkage disequilibrium (r^2^) following Hill & Robertson [48]. In line with the above observations, we observed extended linkage disequilibrium (LD) along the genome in remote Alaskan/Yukon populations (Porcupine and Mile Slough Rivers) and in the Salmon R. population (Thompson R. Watershed, **Fig S2C**). Accordingly, the LD decay - here defined as a r^2^ reaching half of its maximum value - is observed at 25, 12 and 6 kbp in these three populations, contrasting with values around 0.5 kbp observed in the other populations(**Fig S2C**). These results show a large among population variance in effective population sizes, likely associated with the strength of the founder events.

In addition, we constructed a population phylogeny based on a matrix of pairwise *F*_ST_ values computed between each pair of populations(**Fig 2A-B**) and on a matrix of variance-covariance in allele frequencies implemented in Treemix [49] (**Fig S3**). Both analyses confirmed that populations located in the north underwent more genetic drift, in line with the postglacial expansion from the south to the north and the subsequent founder effects associated with upstream colonisation of the rivers from the west to the east.

**Figure 2:**
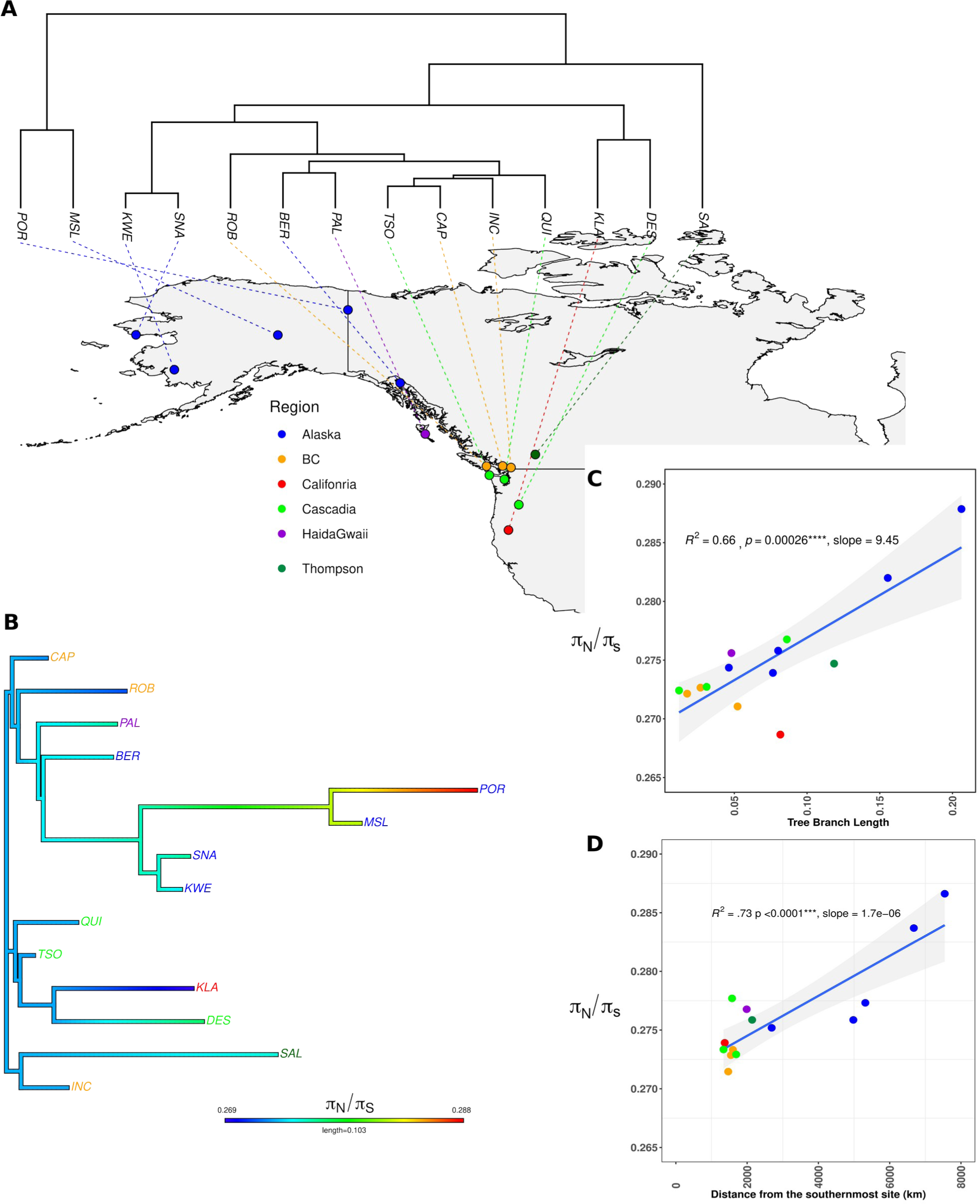
Expansion routes inferred from phylogenetic analyses revealed the extent of the mutation load. A) population phylogenetic tree inferred from the pairwise *F_ST_* values (see also Figure S3) superimposed on the map. Map produced with R using the worldHires data [51]. B) Population specific branches are coloured according to the π_N_/π_S_ values, used as a proxy of the genetic load. C) Correlation between tree branch lengths to the root and the π_N_/π_S_ ratio. D) Correlation between the travelled distance to the southernmost site and the π_N_/π_S_ ratio. The blue line represents the slope of the linear model between the two variables. The grey area in panel C and D represents the 95% confidence interval levels around the regression lines.

To consolidate the above observations, we investigated historical changes in *N_e_* using the Sequentially Markovian Coalescent in SMC++ [50]. This method takes advantage of both the SMC and of the site frequency spectrum to infer change of effective population size though time. We inferred such changed for each population separately (**Fig S2D**), testing a mutation rate of 8e^−9^ and 1.25e^−8^ mutations/bp/generation, which respectively corresponds to the median and the mean mutation rate estimated in Atlantic salmon (*salmo* salar), a closely related salmonid (J. Wang, personal communication). This analysis revealed: *i)* an expansion of most populations approximately 12-20 KyA ago, concomitant with postglacial recolonization, *ii)* a slow and steady demographic decline in the Thompson R. (**Fig S2D**), and *iii)* a split time between all pairwise combinations of populations (median = 16,3 KyA, range = 6,7KyA - 61KyA, **Fig S4**) compatible with the onset of postglacial population expansion and colonisation of different rivers following glacial retreat [41,42]. Using the mean mutation rate yielded similar results with a more recent estimates of split times (median = 9,6 KyA, **Fig S4**) (min = 5 KyA – max = 18 KyA). Overall, SMC++ results indicate that all populations shared a similar demographic history until they began to diverge following the end of the last glacial cycle (see **Note S1** and associated Supplementary Tables).

### Range expansion explained variation in the mutation load

We estimated the mutation load of the populations using different metrics : *i)* π_N_/π_S_*, ii)* count of derived homozygous mutations (including both missense and Loss-of-function (LoF) mutations), and *iii)* total count of derived mutations (including both missense and LoF mutations). We observed a significant positive correlation of the π_N_/π_S_ ratio of each local population and the distance to the southernmost site (see methods), corresponding to the most-likely refugia as discussed in our previous work [42] (R^2^ = 0.73, p<0.0001, **Fig 2D**). Similarly, using the tree branch length to the root of our tree (**Fig 2B**) as a proxy of expansion route revealed a significant correlation with the π_N_/π_S_ ratio (R^2^ = 0.66, p<0.0001, F**ig 2C**). This result was also supported by an analysis considering tree branch lengths inferred with Treemix (**Table S4**). Accordingly, there was a significant correlation between π_N_/π_S_ and two different proxies of the effective population size, namely levels of nucleotide diversity at synonymous sites (π_S_) and SMC++ *N_e,_*, expected to be reduced at the colonization front and in bottlenecked populations (π_S_ in **Fig S5A** and SMC++ *N_e_* in **Fig S5B**). These results support expectations pertaining to variation in mutation load under a model of expansion load (e.g. [28]).

The interpretation of π_N_/π_S_ in populations that are not at mutation-drift equilibrium (say after a bottleneck) is however difficult, since π_N_ will reach an equilibrium value faster than π_S_ because selected alleles will be subject to higher negative selection and undergo a faster turnover [14,52,53]. Thus, the π_N_/π_S_ may not be the best predictor of the total burden of deleterious mutations, as it is potentially affected by demography and selection [14]. To circumvent this limitation, we counted and plotted the distribution of non-synonymous mutations classified according to i) their impact and expected consequences on fitness (*i.e.* missense and LoF mutations) and; ii) segregation patterns (*i.e.* additive (total) load composed of heterozygous and derived homozygous genotypes or recessive (fixed) load composed of homozygous derived genotype) [54]. Just as the π_N_/π_S_, we found a significant association between the derived load and the distance to the southernmost sites both for missense mutations (R^2^ = 0.790, p < 0.001, **Fig 3A**) and for LoF mutations (R^2^ = 0.777, p < 0.001, **Fig 3B**). Similarly, we observed a significant association between the tree branch length to the root and the derived load of missense and LoF mutations (R^2^ = 0.39, p = 0.0096 and R^2^ = 0.40, p = 0.009, respectively, **Table S4**).

**Figure 3:**
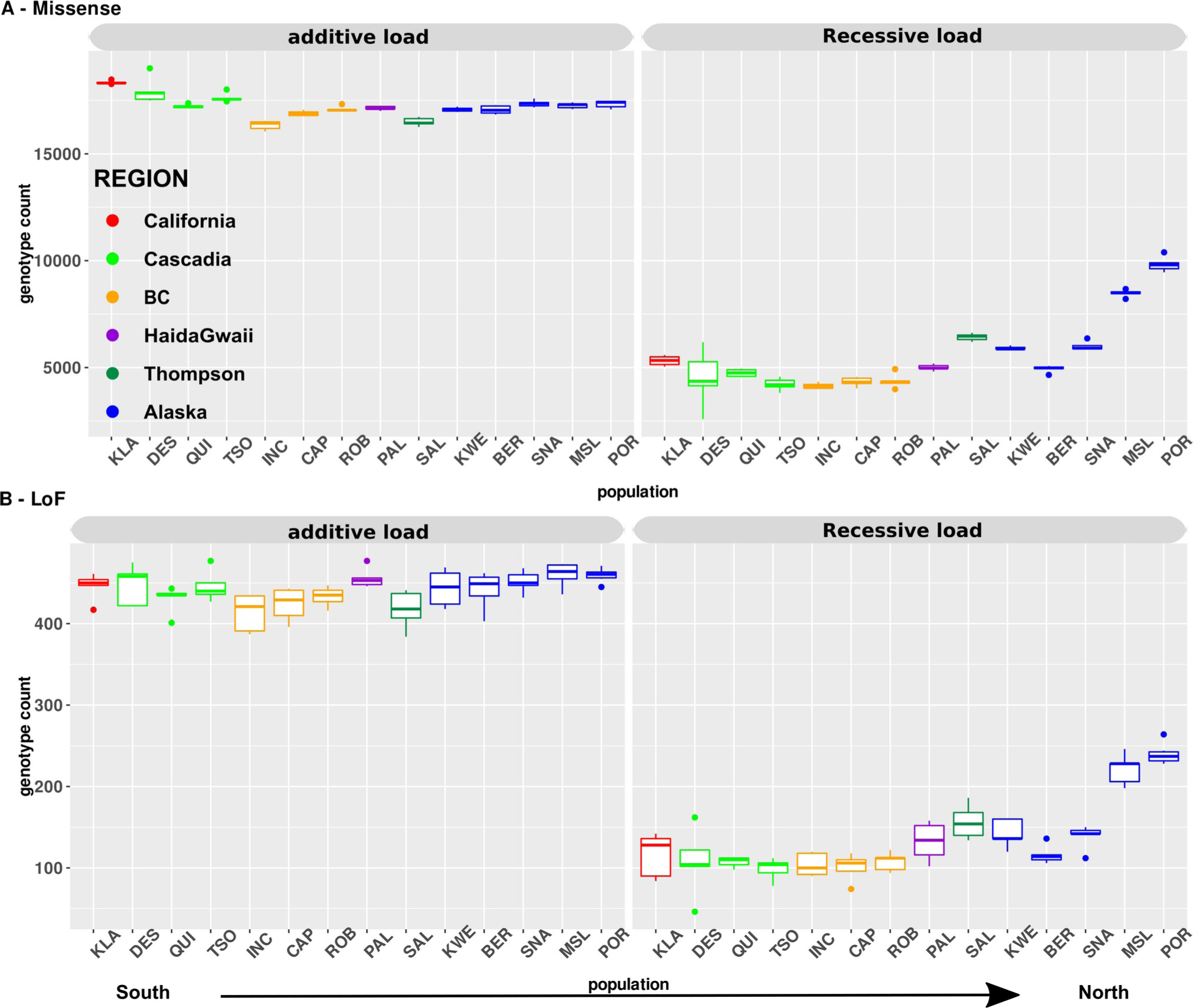
Number of deleterious alleles per riverper individual for all populations sorted from the South to the North. *The plot* shows the additive load (left panel) and recessive derived homozygous load (right panel) for A) missense mutations and B) Loss-Of-Function (LoF) mutations. No strong differences are observed in the additive load among populations. In contrast, significant differences were observed for the recessive load in populations at the expansion front both for missense mutations and LoF mutations and this was significantly correlated with the distance to the southernmost site. Each color represents a major regional group. Abbreviation for each site is provided in Table S01.

#### The most deleterious mutations are efficiently purged across the range

The total number of missense mutations was significantly more abundant in the more southern populations (*i.e* California, Cascadia, **Table S5** for all Tukey-HSD *p-values*) than in northern ones (**Fig 3A**), while it was nearly constant from the south to the north regarding LoF mutations (**Fig 3B**), with no significant differences among populations (**Table S5**). The southernmost populations displayed a higher load of mutations in heterozygous state (**Fig 3**), as expected due to their higher historical effective population size, favouring the segregation of recessive mutations hidden in a heterozygous state [42,55].

#### The fixed load increases with population expansion

The fixed load (i.e. count of derived homozygous sites) increased from south to north for both missense and LoF with the most extremes samples, including Mile Slough (MSL), Porcupine (POR) and Salmon (SAL) rivers being the most significantly loaded (**Table S5**), as expected due to founder events and allele surfing for Alaskan/Yukon samples and due to bottlenecks for the Salmon River (**Fig 3** right panel).

#### Range expansion have reduced the efficacy of selection at range margin

We predicted that selection efficacy will be reduced at the expansion front and in populations with a lower *N_e_.* To test this, we measured three parameters, namely the rate of non-adaptive and adaptive amino-acid substitutions relative to neutral divergence (ω_NA_ and ω_A_, respectively; with ω_A_ = d_N_/d_S_ - ω_NA_) and the proportion of non-synonymous amino-acid substitution that result from positive selection (α = ω_A_/(d_N_/d_S_)) using the software Grapes [56] using maximum likelihood estimation under a population genetic model. A general expectation is that populations at the expansion margin will exhibit a higher rate of non-adaptive substitutions (ω_NA_), a lower rate of adaptive substitutions (ω_A_) and a lower proportion of amino-acid substitutions (α).

We observed a reduced efficacy of purifying selection likely associated with the multiple founder events and bottlenecks which resulted in a decrease rate of adaptive substitutions (ω_A,_ R^2^ = 0.37, p = 0.013) and increased ω_NA_ as a function of the distance to the South (R^2^ = 0.57, p = 0.0011, **Fig 4A**). We also found a significant positive correlation between ω_NA_ and the distance to the ocean (R^2^ = 0.52, p = 0.002, **Fig S6A**) and conversely a negative correlation between ω_A_ and the distance to the ocean (R^2^ = 0.40, p = 0.00895, **Fig S6B**), suggesting that upstream populations displays a lower adaptive potential. This observation thus suggests that the population expansion had impacts on both the adaptive and non-adaptive substitution rates.

**Figure 4:**
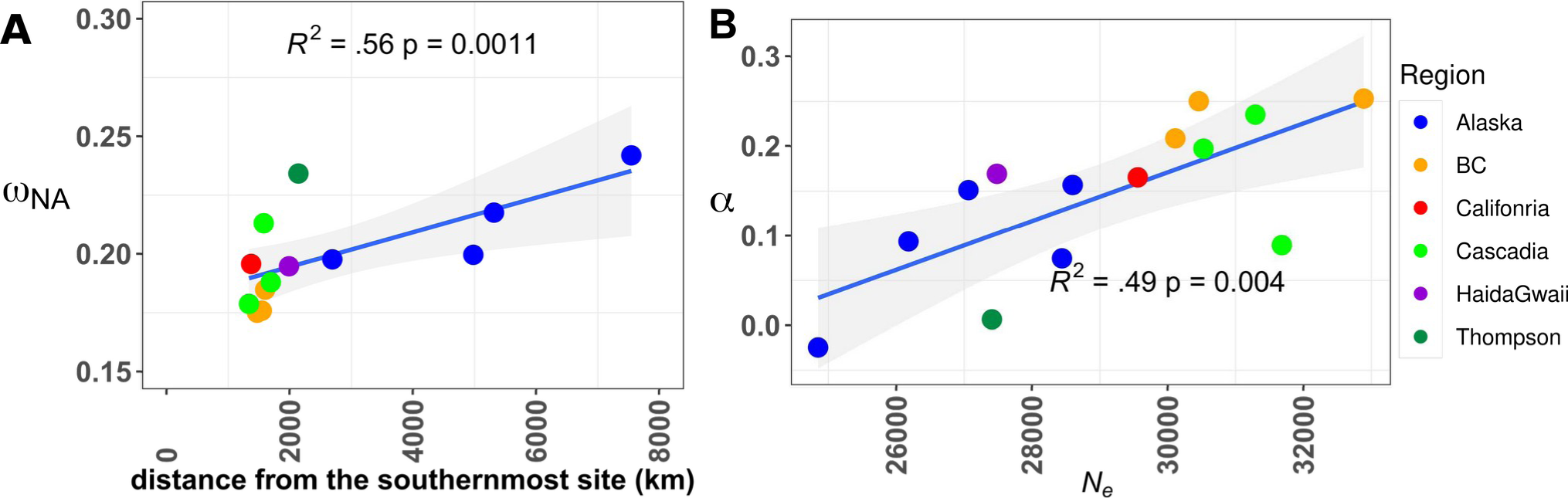
Efficacy of selection decreases at the expansion front and as a function of long-term effective population size. A) increased rate of nonsynonymous non-adaptive substitutions (ω_NA_) in northernmost populations of Coho salmon (Alaska) and in the bottlenecked Thompson River. B) Correlation between α and historical variation in the coalescent effective population size estimated from SMC++. See also **Fig S7-S8** for correlations based on tree branch lengths. Note that the distance from the southernmost site was computed considering a Californian sample, from a previous study [42], for which no WGS data were available.

Similarly strong correlations were observed when considering the tree branch length to the root (details in **Fig S7-S8, Table S4**). The Salmon R. population (Thompson R. watershed), which suffered a recent decline in abundance, supposedly due to the sustained release of hatchery fish derived from few individuals and the Ryman-Laikre effect [57] (see discussion below), also displayed a high ω_NA_.

We also predicted a higher α value in populations with higher *N_e_* and, conversely, a decreased value of α in populations with lower *N_e_.* We observed a strong positive correlation between α and the synonymous nucleotide diversity (π_S_) of each local population, which represents a good predictor of the long term (coalescent) *N_e_* (R^2^ = 0.63, p = 0.0004, **Fig S9**). To more directly test the association with *N_e_*, we correlated SMC++ based estimates of *N_e_* (averaged over a 200KyA window) with α and recovered significant correlations (R^2^ = 0.49, p = 0.004, **Fig 4B**). It is noteworthy that a good correlation between proxies of *N_e_* obtained from SMC++ and π_S_ was also observed (R^2^ = 0.68, p = 0.00017, results not shown). Altogether, the above results provide empirical support pertaining to the evolutionary consequences of allele surfing at expanding range margins, in particular regarding the loss of adaptive potential and the mutation burden.

#### Recombination rates shape the deleterious mutational landscape

In addition to the spatial structure associated with the postglacial recolonization, we investigated the genome-wide variation in mutation load. Such variation could be associated with the occurrence of structural variants (e.g. chromosomal inversions) which may incur a significant load because deleterious recessive mutations may freely accumulate in the absence of recombination, as observed for instance in sex chromosome and related supergene-like architecture [58]. To test this hypothesis, we used the GC content at third codon position (GC3 hereafter) as a proxy of the rate of recombination [59,60] (see Sup. Note S2 for an explanation). We observed strong correlations between levels of GC3 and π_N_/π_S_ ratio for all Coho salmon populations (R^2^ range = 0.938 – 0.955, p<0.001) except for population MSL (R^2^ = 0.252; p = 0.0038) (**Fig 5A**). An analysis focused on GC-conservative sites, which are not affected by GC-biased Gene Conversion (gBGC), revealed similarly strong patterns across all Coho salmon populations (**Fig 5B**, R^2^ range = 0.665 - 0.876, p < 0.01). π_N_/π_S_ ratios estimated based on all sites or GC-conservative-sites only are highly correlated (Pearson r = 0.909, p <0.001). We tested the generality of this relationship using four other closely related Pacific salmon species and found strikingly similar pattern in Chinook salmon (R^2^ GC3 ∼ π_N_/π_S_ = 0.9338, p = 1e-11) Sockeye and Kokanee salmon (R^2^ range = 0.81 - 0.96, p <0.001), as well as Rainbow trout populations (R^2^ range = 0.95 - 0.96, p <0.001, **Fig 5C**).

**Figure 5).**
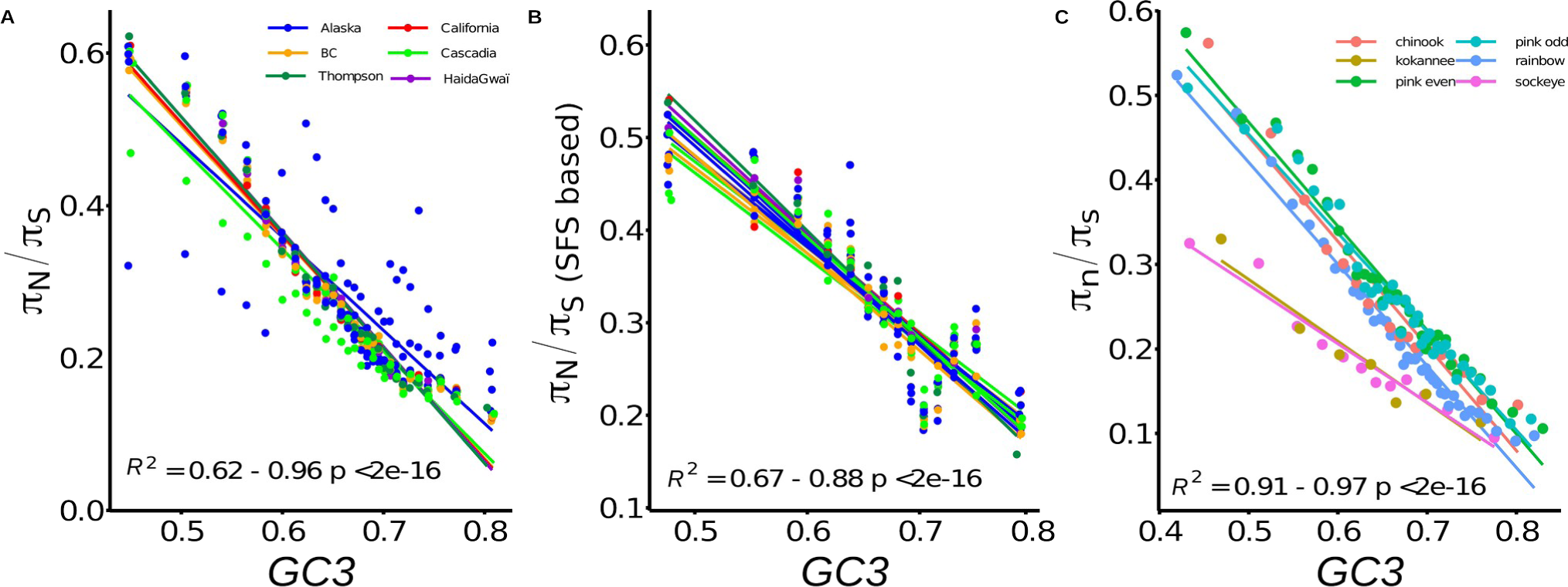
The deleterious load is determined by variation in GC3 content in multiple salmonids. A) Correlations between the deleterious load (π_N_/π_S_) and GC content at third codon position in multiple Coho salmon populations. Each line represents the values colored by major regional groups (See **Fig S10A** for detail). B) Correlations between the deleterious load (π_N_/π_S_) and the GC content at third codon position considering an independent method based on the site frequency spectrum at GC-conservative sites. C) Correlation between the deleterious load (π_N_/π_S_) and GC content at third codon position in sockeye salmon ecotypes (sockeye and kokanee), chinook, pink salmon and in rainbow trout. Averages are provided for rainbow trout, salmon and kokanee at the species level, see **Fig S10B-C** for detail by population.

We then analyzed the difference in the correlations between levels of GC3 and π_N_/π_S_ ratio among all Coho salmon populations. Indeed, the intensity of this correlation highlights the effect of recombination rate variation on the efficiency of purifying selection [61]. The populations at the expansion front (Mile Slough R., Porcupine R., Snake R.) in Alaska exhibit the lowest correlations between GC3 and π_N_/π_S_. If both demographic factors (*i.e.* distance to the source here) and genomic factors (*i.e.* recombination) interact to shape the load, we may predict a positive relationship between these two factors because the northernmost populations should have an even higher load in regions of low recombination. We tested this hypothesis using a linear model between the slope of the regression between π_N_/π_S_ and GC3 and the distance to the ancestral source populations did not reveal such a relationship (p = 0.133, **Fig S11**).

#### Regions of residual tetraploidy revealed drivers of the load in Coho salmon

We tested the hypothesis that regions with residual tetraploidy exhibit a reduced load due to increased efficacy of purifying selection due to higher population sizes (4*Ne* rather 2*Ne*). Contrary to this expectation, increased deleterious load was observed in regions of residual tetraploidy (3,700 genes) as compared to diploid regions (**Fig 6A**, red dotted line mean π_N_/π_S_ > 0.35, diploid region π_N_/π_S_ < 0.30). Given that we also observed lower levels of recombination in regions with residual tetraploidy compared to re-diploidized genomic regions (**Fig S12,** p < 0.0001, W Mann-Whitney = 3.04e^7^, see also **Fig S13** and **table S6** for differences in recombination among populations), our results suggest that this higher π_N_/π_S_ could be mostly due to lower recombination rates. Another expected genomic consequence of this lower recombination rate is a higher load of transposable elements [62]. When computing the relative length of TE, *i.e.* the length of TE corrected by the chromosome length (see methods), we found a significant enrichment of TE in the regions of residual tetraploidy as compared to diploid chromosomes (**Fig 6B**, p <0.0001, WMann-Whitney = 1.36e^10^). This tendency was also observed across the different TE categories (**Fig S14**). To more directly test the *N_e_*-effect hypothesis, we eliminated the effect of the recombination rate by comparing the load across similar bins of GC in diploid vs. tetraploid regions. The π_N_/π_S_ was systematically higher in diploid regions than in the tetraploid ones after excluding the class with the lowest GC content (**Table S7, Fig 6C**) indicating that the load was significantly higher in diploid compared to tetraploid regions, with the notable exception of regions with extremely reduced recombination. Therefore, it is still possible that increased efficacy of selection is at play in recombining regions of residual tetraploidy, following the hypothesis of higher effective size in these regions.

**Figure 6:**
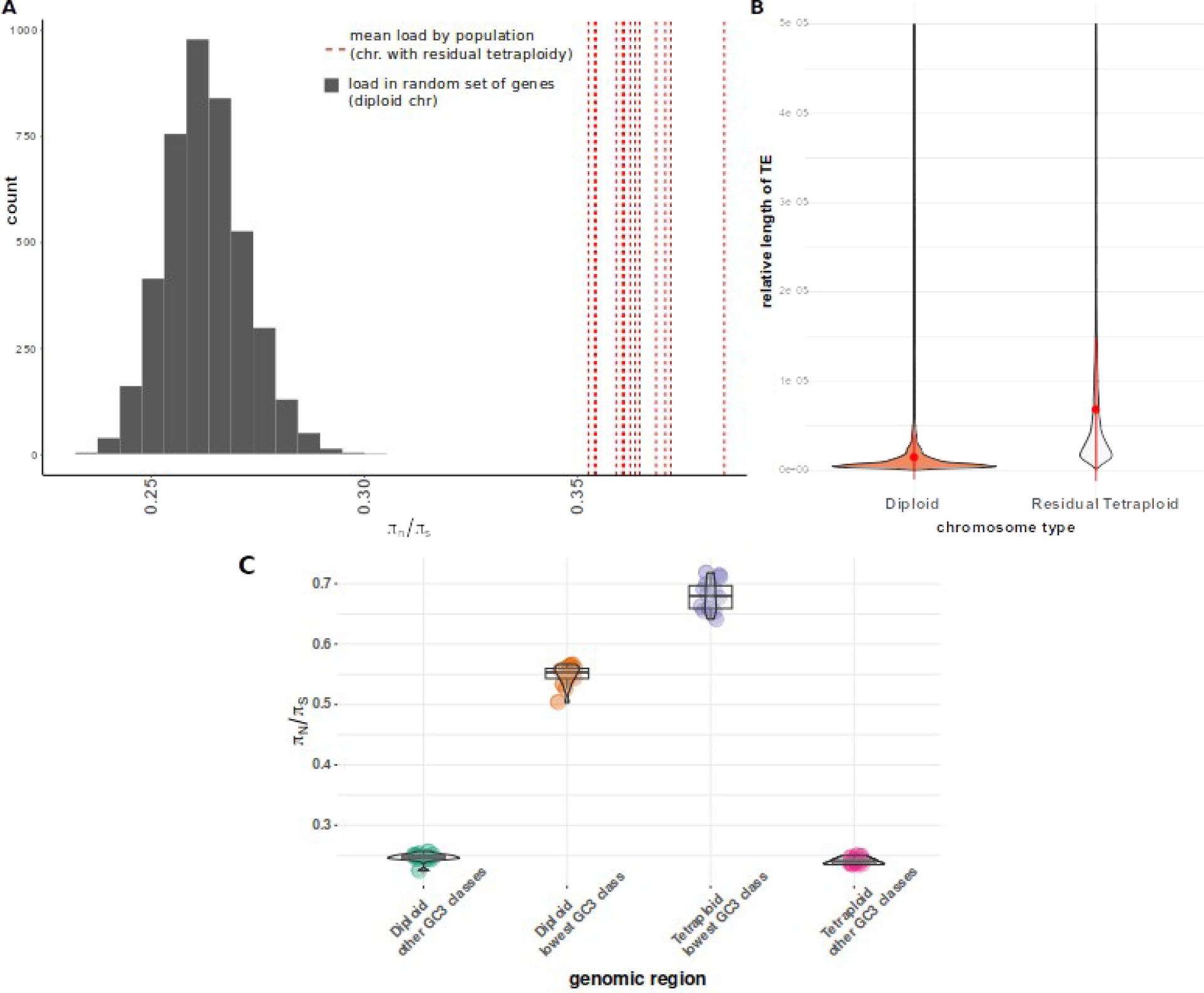
Increased load in region of residual tetraploidy. **A**) Distribution of π_N_/π_S_ ratio when considering all genes in regions of residual tetraploidy (red lines) for each population compared to a set of 200 randomly generated samples of 4,000 genes in the rest of the genome (gray histogram). **B)** Violin plot showing that regions of residual tetraploidy displayed significantly longer transposable elements when compared to diploid regions. Orange = diploid chromosome. Gray = chromosome with residual tetraploidy. Red point = mean +/− 1*sd. **C**) Difference in the distribution of πN/πS ratio in different genomic regions of the Coho salmon. Each dot represents the value observed for a given population in either diploid or tetraploid region with the “lowest” GC3 value (corresponding to low recombination) compared to the rest of GC3 values (labelle “other GC3”, corresponding to intermediate to high recombination). The GC values were computed in 4mb windows separately for diploid and tetraploid regions.

## Conclusion

The role of genetic drift, recombination, selection and variation in inheritance in affecting the deleterious load and efficacy of natural selection is a central question with fundamental and applied consequences for biodiversity.

Using population genomics analyses, we investigated the evolutionary consequences of recent demographic events in Coho salmon. Our results supported gene surfing across North America [42]. Results also supported lower *N_e_* at the northern range expansion front, which induced two main evolutionary consequences: a surf of slightly deleterious mutations and a putative reduction of the adaptive potential, as expected under the nearly neutral hypothesis of molecular evolution [12].

We further demonstrated that one population, the Salmon River (Thompson R. watershed) displayed an increased fixed load and decreased selection efficacy compared to southern populations. Previous studies showed that this population is genetically isolated from others [42,43,63] and displayed genomic footprints of a bottlenecked population such as long runs of homozygosity [41]. This population has been subjected to extensive hatchery enhancement (from a single population) to circumvent the decline of Coho salmon in the Thompson drainage [64]. From a theoretical standpoint, if a few captive individuals were used as parents for subsequent releases, then an increase in inbreeding and a reduction in effective population size are expected, a phenomenon called the Ryman-Laikre effect [57] [65]. Consequently it is possible that the increased π_N_/π_S_, ω_NA_, and fixed deleterious load could be explained by both long-term evolution at low *N_e_* and by the Ryman-Laikre effect associated with hatchery enhancements. Regardless, this suggests that careful enhancement needs to be performed with a diversity of parents to maximize genetic diversity of supplemented populations. This result has implications for fisheries enhancement, as well as genetic rescue programs aiming at reducing the inbreeding load of declining populations and restoring the fitness of these populations.

It is noteworthy that the southernmost population (Klamath R, California) was not characterized by the highest α value. Several non-exclusive hypotheses may explain this pattern: first, some Californian populations are known to have undergone strong recent (human-driven) decrease in abundance due to both habitat degradation and climate change, which may leave a detectable footprint in the genome [66]. The recent demography could have an impact on the accuracy of the fit of the synonymous and nonsynonymous SFS under Grapes. Second, the Klamath R. population is located in the far upstream part of that river, and may have undergone a founder event when reaching this upper part, a hypothesis that is supported by the correlations we observed between several metrics and distance to the river mouth. Finally, we did not include the southernmost populations from California, which are more likely to be more ancestral, as we previously reported [42].

Since the use of π_N_/π_S_ ratio can be criticized [14,67,68], especially when used to quantify the load within species, we computed additional metrics [56] to more directly estimate mutation load, as advocated by others [14,69]. In this case, we found small differences in the additive load among Coho salmon populations, but a linear increase in the recessive load as a function of the distance to the southernmost sites and as a function of the tree branch length, both are used as proxies of the expansion route.

We demonstrated that the π_N_/π_S_ ratio was negatively correlated with the GC content at third codon position, which represents a good proxy of the local recombination rate (see **note S2**). The negative correlation between π_N_/π_S_ and GC3 was repeatedly observed across different salmonid species using both a sequence based estimate of π_N_/π_S_ and an estimate based on GC-conservative site (non-affected by gBGC) [60]. These results indicated that recombination plays a key role in explaining the variation in the mutation load along the genome in salmonids. Our study empirically supports theoretical work about the accumulation of slightly deleterious mutations in non-recombining regions [30,32]. Similarly, increased prevalence of deleterious mutations in low recombining regions have been reported in plants [70] and in human populations [71–73]. In particular, recent work in human populations have shown that both variation in demographic history (i.e. change in effective population size) and recombination rates are affecting allele-specific expression of harmful mutations [71]. The authors showed that allele specific expression causes underexpression of harmful mutations more efficiently in normally and highly recombining regions compared to low recombining regions. They further documented variation of this process among populations with varying demographic histories.

In line with the key role of recombination rate, we observed that only regions of residual tetraploidy with extremely low recombination rate (lower GC content) displayed an increased load in Coho salmon, accumulating more transposable elements, suggesting efficient selection in more “normally” recombining region of residual tetraploidy. In particular, our results indicate that i) the regions of residual tetraploidy (4Ne) with a normal recombination rate do not display an increased load, which may be expected under the nearly neutral theory if higher Ne is associated with the efficacy of selection; ii) only region of extremely low recombination rate (lower GC content) displays the highest load, highlighting once again the primary role of recombination in shaping the load, with a higher contribution than Ne. A third factor is the variation in dominance of deleterious mutation. Indeed, a recent simulation study found that in the case of hard sweeps, dominance of recessive mutation was a central aspect determining the signal of sweeps in polyploid genomes [39]. A detailed investigation of this aspect in regions of residual tetraploidy was beyond the scope of our study but would be worthy of further investigation.

The detailed consequences of ongoing rediploidization have been extensively studied at the regulatory levels elsewhere, with support for an increased load (higher d_N_/d_S_ and TE load) in duplicated genes undergoing lower expression [74]. Recent studies in polyploid plant species have also documented various evolutionary consequences of such duplication on local variation in the load and efficacy of selection [35]. F or instance in *A. arenosa,* a reduced efficacy of purifying selection was suggested in tetraploids (4X) genome compared to diploids (2X), and the authors suggest that this is because deleterious alleles are better masked in autotetraploids [38]. Similarly, a recent work in the allopolyploid cotton *(Gossypium)* demonstrated that this species accumulates more deleterious mutations than the diploid species [75]. This observation supports a theory proposed by Haldane that recessive deleterious mutations accumulate faster in allopolyploids because of the masking effect of duplicated genes [76]. To sum up, our results concur with predictions from the nearly neutral theory of molecular evolution [12], in which slightly deleterious mutations are effectively neutral and purged effectively in regions of higher recombination, except perhaps in populations at the extreme of the expansion front. Interestingly, these results indicate that regions of low-recombination re-diploidize later than other genomic regions. We are not aware of any paper having documented such patterns so far. Further studies of this process across more species would be welcome to validate the generality of this observation.

Finally, our results have implications for conservation practices. We showed that the additive load is approximately constant, indicative of efficient purging across populations for both missense and LoF, similar to some recent studies on several plant and animal models [77–80]. However, results indicate that population at higher latitude from the Yukon watershed (Mile Slough R., Porcupine R.) or from the bottlenecked Salmon R. have not entirely purged the most deleterious mutations, including missense and LoF mutations, which may impose a fitness cost to these populations. Further empirical evidence for a causal link between the putative fitness cost of LoF mutations and adaptive phenotypic variation will be necessary to validate our observations. With this caveat in mind, our results for the Salmon R could guide practices in supplementation programs. For instance, choosing a diversity of parents from moderately differentiated populations of modest size may help increase the levels of heterozygosity and mask the expression of recessive deleterious mutations [55]. This strategy could reduce the occurrence of deleterious alleles and counteract the Ryman-Laikre effect described above. Moreover, populations (e.g. most upstream Alaskan/Yukon populations) for which our results suggest a reduced adaptive potential are also the most strongly exposed to rapid climate change, as the rate of temperature increase is most rapid at higher latitudes [81]. In all cases, maximizing the connectivity among populations and limiting habitat degradation appears as fundamental strategies to maintain high effective population size and increase the adaptive potential of Coho salmon to the multiple ongoing anthropogenic pressures [82]. More generally, how best to manage declining populations and guide conservation policies in these conditions to minimize the load and/or maximize genetic diversity is another debated issue [55,83–85]. In the meantime, while conservation genomics undoubtedly has a major role for the short-term preservation of endangered species, this should not override the crucial need for reducing human impacts on natural ecosystems to preserve biodiversity over long time scales [86].

## Methods

### Sampling design for Coho salmon

We sampled and sequenced 71 individuals representing 14 populations distributed from California to Alaska (**Table S1**). A set of 55 individuals was sequenced on an Illumina HiSeq2500 platform [38] and the other 16 individuals were sequenced on a NovaSeq6000 S4 platform using paired-end 150 bp reads. Reads were processed using fastp for trimming [87], and mapped to the most recent Coho reference genome (https://www.ncbi.nlm.nih.gov/assembly/GCF_002021735.2/) using bwa mem v2 [88]. Reads with a minimum quality below 20 were discarded with samtools v1.7. Duplicates were removed with picard (http://broadinstitute.github.io/picard/). SNP calling was performed using GATK v4.2 [89] using our pipeline available at github.com/quentinrougemont/gatk_haplotype/. We generated a Haplotype gVCF for each sample individually, combined all gVCF and then performed a joint genotyping. We checked the variants quality score of our data and filtered our genotypes according to their quality following GATK best practices and based on quantiles distributions of quality metrics. We excluded all sites that did not match the following criterion: MQ < 30, QD < 2, FS > 60, MQRankSum < −20, ReadPosRankSum < 10, ReadPosRankSum > 10. We also excluded multiallelic SNPs, as well as indels. Genotypes with a depth lower than 6 or higher than 100 reads were also excluded to remove low confidence genotypes potentially associated with paralogs. Finally, we also generated a separate vcf file using the --all-sites option to obtain a file with invariant position to reconstruct sequence data (see the **Genetic load estimation in Coho salmon and related species** section). A total of 14,701,439 SNPs were identified without missing data. Population structure was evaluated using a principal component analysis (PCA) performed using Ade4 [90] on a set of LD-pruned SNPs without missing data (1,739,037 SNPs) identified stringently with plink1.9 [91] (command indep-pairwise 100 50 0.1).

#### Outgroup dataset

In order to test the generality of the relationship observed between π_N_/π_S_ and GC3 we took advantage of the newly assembled reference genomes and resequencing data from other closely related Pacific salmonid species with similar demographic histories[92–94]. Sockeye salmon published by [92] were retrieved from NCBI PRJNA530256. Samples with a high number of individuals per ecotype (Sockeye and Kokanee) were chosen from the NCBI table. We retained a total of 5 Kokanee populations and 5 Sockeye populations (from Fraser & Columbia watershed described in T**able S1 and Table S3**). Three Chinook salmon samples were provided by B. Koop (also available at NCBI PRJNA694998). Additionally, 11 samples of “even” and 10 samples of “odd” pink salmon (*O. gorbuscha*) were downloaded from PRJNA 556728 and included in the analysis. Here “even” and “odd” refers to Salmon returning to their natal rivers in “even” and “odd” years to spawn, leading to a temporal isolation of these ecotypes [93]. These were indeed clearly separated based on a PCA. Finally, a number of rainbow trout available from NCBI PRJNA386519 were used (n = 19 from 3 random populations showing genomic differentiation based on a PCA).

Each sample was downloaded and mapped onto its species’ reference genome downloaded from NCBI and using the exact same procedure as described above relying on fastp, bwa-mem2, picard and GATK 4.2.5.0 to generate a final vcf filter based on usual GATK quality criteria and variance in sequencing depth. For each species, we then quantified the load using the π_N_/π_s_ ratio with the procedure described below for Coho salmon. For the Sockeye/Kokanee ecotypes, the Sockeye is a fully migratory ecotype, whereas the Kokanee is a resident (non-migratory) ecotype that typically comprises more isolated populations. The two alternative ecotypes are sometimes found in similar locations, with three rivers from our sampling design including both ecotypes (Table S03). For all other species we tested the relationship between π_N_/π_S_ and GC3 (see below).

### Ancestral and derived alleles identification

To accurately recover the ancestral allelic states, we used three outgroup species; including the chinook salmon and rainbow trout sample (see above, n = 5 for rainbow trout) plus data from Atlantic salmon (n = 5, SRP059652). Each individual was aligned against the Coho salmon V2 genome (GCF_002021745.2) using GATK with the same procedure as above and calling every SNP using the –all-site mode to obtain invariant positions. We then determined the ancestral state of each SNP if 1) the SNP was homozygous in at least 90% of the individuals from our three outgroups, and 2) matched one of the two alleles identified in Coho salmon. Otherwise, the site was inferred as missing and was not used in subsequent analyses of the load. In addition, we reconstructed a consensus ancestral fasta sequence using the doFasta option from angsd [95]. This was used for demographic reconstruction detailed in **Note S1.**

### Demographic reconstruction

We first tested our prediction that genetic diversity (observed heterozygosity *H_o_*) decreases towards the North following our «out of Cascadia» model previously inferred [42,43]. Conversely, we verified that the *β*_ST_ coefficient, a measure of both genetic differentiation and ancestrality [46] increases northwards the North, as expected due to isolation by distance. *β*_ST_ and observed heterozygosity (H_o_) were measured using the hierfstat R package [96] Details about β_ST_ computation are provided in Weir & Goudet [46] and H_o_ was computed following Nei (1987)[97] as 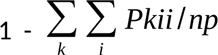 with *Pkii* the proportion of homozygotes *i* in sample *k* and *np* the number of samples. Oceanic coastal distances were computed using the marmap package [98] and waterway distance was computed using ArcGIS. Then we summed the 2 distances to obtain the distance to the most likely refugia that we identified in our previous studies [42,43]. A shapefile of the rivers used here is available at https://github.com/QuentinRougemont/selection_efficacy/tree/main/00.raw_data. Under postglacial expansion from a single refugium, the general hypothesis would be that all sampled populations should follow a common temporal trajectory of a population decline (bottleneck due to founder events by few individuals) followed by a (strong) increase in *N_e_*corresponding to the expansion phase. To test this hypothesis, we inferred temporal changes in *N_e_* using SMC++ [50]. SMC++ works similarly to the PSMC model but takes into account the Site Frequency Spectrum (SFS) and is better suited to large sample sizes. Estimates of changes in population size were performed for all populations. To validate the fact that expansions are indeed postglacial, splitting time was estimated between all pairs of samples from different geographic areas based on the joint SFS (n = 75 pairwise comparisons). A generation time of 3.5 years, well documented for Coho, and a mutation rate of 8e^−9^ mutation/bp/generation were applied (corresponding to the median substitution rate inferred in Atlantic salmon, J. Wang, Personal communication). We also compared the results to the mean substitution rate of 1.25e^−8^ mutation/bp/generation also inferred by Wang.

In addition, pairwise linkage disequilibrium provides valuable information regarding population size and inbreeding. We computed the squared correlation coefficient between genotypes in each sample and all populations separately using popLDdecay [99]. We used a MAF of 5% in each population, keeping between 3.7 and 6.4 million SNPs with populations having undergone stronger bottleneck/founding events displaying the lowest amount of variation. We estimated LD decay by plotting LD (R ^2^) against physical distance measured in base pairs in ggplot2 [100] package in R.

We observed slight discrepancies between our SMC++ estimates of divergence time and our previous work based on the site frequency spectrum [42]. To investigate this, we performed a new set of inference based on the unfolded joint site frequency spectrum (jSFS) using ∂a∂i [101] and a new set of refined models as detailed in **supp. Note S1, Table S8-S9**.

To reconstruct the population expansion routes we computed Weir and Cockerham [102] *F*_ST_ values among populations using Vcftools [103] and constructed a phylogenetic population tree using the R package ape [104]. We then computed the tree branch lengths to the root using the function distroot from adephylo [105]. We also explored broad relationships among populations with treemix [106]. We fitted a model with an increasing number of migration edges. We chose a model with K = 3 migration edges as all edges displayed significant p-value and captured a high proportion of explained variance. Fitting more edges decreased the p-value without really improving the fit to our data. Similarly, we computed the tree branch length to the root using the function adephylo and compared our results to those from *F*_ST_-based values. The resulting tree branch length values were used as a proxy for the expansion routes, and we tested their correlation using linear models in R, with all our metrics of deleterious load (π_N_/π_S_, total number of putative homozygous derived deleterious alleles (recessive load) and total number of deleterious alleles (both in homozygous and heterozygous states, additive load) for derived missense and LoF mutations) and metrics of selection efficacy (ω_NA_, ω_A_, α detailed below). Similar inference was performed, but using the reconstructed distances to the southernmost site included in our previous studies [42,43]. This distance was computed by summing the oceanographic distance and river distances described above.

### Population-scaled recombination rate

Statistical phasing of the Coho whole genome sequences was performed using the Shapeit software [107], considering all individuals at once. We then estimated effective recombination rates (ρ=4.*N_e_*.r where r represents the recombination rate per generation and *N_e_* is the effective population size) along the genome using LDHat [108]. Phased genotypes were converted into LDHat format using vcftools after excluding SNPs with MAF < 10% since rare variants are not informative for such inferences. Following the guidelines of the LDHat manual, the genome was split into fragments of 2,000 SNPs with overlapping windows of 500 SNPs. We measured recombination rates independently for each population. Differences in the distribution of population-scaled recombination was visualized using violin plot (**Fig S13**) and statistically tested using Tuckey HSD tests in R (**Table S5**).

#### Genetic load estimation

##### π_N_/π_S_ estimates

The approach developed in [59] was used to reconstruct fasta sequences for each individual. For each species (Coho and the other salmonids), coding sequences (CDS) from each reconstructed sequence were extracted using the gff files available with the reference genome to estimate the nucleotide diversity (π). We also concatenated the CDS sequences into different classes according to their length and computed π_N_ and π_S_ over 4-Mb concatenated gene space windows. Such large windows reduce the stochasticity due to the low π_S_ values in Coho salmon.

##### Identifying potential deleterious non-synonymous alleles

We tested the difference in count of non-synonymous mutations in each local population of Coho salmon across non-synonymous missense mutations (putatively deleterious) and Loss of Function (LoF) mutations (likely to be strongly deleterious) identified with SNPeff. We analyzed data in two ways: first, we counted the total number of putative homozygous derived deleterious alleles (recessive load) per individual as well as the total number of deleterious alleles (both in homozygous and heterozygous states, additive load) using: Ntotal = 2 Χ N_homo_ + N_hetero_ [28]. These individual values were then averaged per population. We tested for differences in the distribution of these counts among populations using an ANOVA followed by a TukeyHSD test. The p-values were corrected using a Bonferroni correction. We also computed mean derived allele frequencies (DAF) in all sampling locations and across mutation categories (synonymous, non-synonymous missense and LoF). We also applied the commonly used Provean [109] software based on a random set of non-synonymous mutations (but see **Note S4** for a brief discussion regarding the limitations).These results were then compared with results from non-synonymous mutations (**Fig S15**).

##### Correlation between π_N_/π_S_ and recombination

We computed the GC content at third-codon positions of protein coding genes (GC3) which has been shown to be an accurate proxy of local recombination rate in other species and these positions are generally silent [85,86]. To compute π_N_/π_S_ values we sorted genes by ascending GC3 values, which enabled us to obtain a ratio based on genes with similar GC3 values. Moreover, we also used the site frequency-based approach proposed by Rousselle et al. [60] to estimate the π_N_/π_S_ ratios. This approach enabled us to compute SFS separately for GC conservative sites (A<->T and C<->G mutations), that is, not affected by GC-biased Gene Conversion (gBGC).

Finally, we measured the correlation between GC3 and π_N_/π_S_ using linear models. We replicated these analyses considering only genes (n = 3,500) and SNPs in regions of residual tetraploidy (8% of the genome).

##### DFE estimation and rate of adaptation in Coho salmon

We estimated the rate of non-adaptive and adaptive synonymous substitutions (ω_NA_ and ω_A_, respectively; with ω_A_ = *d_N_*/*d_S_* - ω_NA_) and the proportion of amino-acid substitution potentially resulting from positive selection (α = ω_A_/(*d_N_*/*d_S_*), with d_N_ being the rate of non-synonymous substitutions, d_S_ being the rate of synonymous substitutions). To do so, we used the method implemented in Grapes v1.0 which builds upon the approach of [110]. Grapes models the effect of favorable mutations on the non-synonymous site frequency spectrum (SFS) while accounting for various confounding factors distorting the SFS (demographic change, linked selection, genotyping errors, SNP misorientation).

We used the following parameters in Grapes v1.0: we assumed a negative Gamma distribution to the synonymous and non-synoymous SFS (parameters GammaExpo in Grapes and parameter “unfolded” site frequency spectrum). To obtain suitable data, we converted the quality filtered whole genome vcf file into a fasta file for each population containing sequence information for each individual in the population (using vcf2fasta available at https://github.com/QuentinRougemont/selection_efficacy/tree/main/08.piNpiS/). This file was then converted into a site-frequency spectrum using bppml [111]. We required at least 10 sites without gap to keep a given site (-gapN_site), with a maximum proportion of gap of 0.5 (-gapN_seq), a minimum number of complete codon of 6 (-min_nb_codon), we did not allow frameshift (remove_frameshift parameter), we also required a sample size of 8 (-sample_size), the number of initial/terminal position in which Stop codons or FrameShift codons are tolerated was set to 20 (-tolerance_zone), we did not allow Stop or FrameShift codons between the two tolerance zones (-allow_internal_bc parameter). The kappa value (i.e Ts/Tv ratio) was set to 1.6 and we used an unfolded SFS using the three species as an outgroup. Fitted parameters of the DFE were used to compute the expected d _N_/d_S_ under near neutrality, which was compared to the observed d_N_/d_S_ to estimate α, ω_NA,_ ωa.

#### Differences in load for region of residual tetraploidy in Coho salmon

##### π_N_/π_S_ comparison

In the Coho reference genome, Chromosomes 1 to 30 are considered to represent generally diploid chromosomes, whereas chromosomes 31 to 38 represent those with a clear signal of residual tetraploid [41]. We took advantage of this specificity regarding chromosome evolution to contrast the load for the 3700 genes in regions of residual tetraploidy (averaged across all genes for each population). For diploid regions, we generated 200 datasets of 4,000 genes randomly sampled and then estimated the load for each of these datasets.

##### TE annotations

We used the TEs annotation file from repeatmasker [112] (made available on NCBI for the reference genome [41]) and tested for difference in the length of TEs between diploid and tetraploid regions, after correcting for the difference in chromosome length.

## Data availability

New sequencing data are currently being deposited on NCBI SRA under project PRJNA982773. The coho vcf is available online at 10.5281/zenodo.8027705. All scripts to reproduce our analyses are currently being deposited online at https://github.com/QuentinRougemont/selection_efficacy.

## Acknowledgements

We are grateful to Alysse Perreault-Payette and Bérénice Bougas for help in DNA extraction of additional individuals for whole genome sequencing. We thank Benoit Nabholz for the initial development of the π_N_/π_S_ C++ app we used here. We are grateful to the genomic platform at Compute Canada (Canada), IBIS (Canada), and MBB (Montpellier, France) for providing computational capacity as well as Bird (Nantes, France) for data storage. This research was carried out in conjunction with EPIC4 (Enhanced Production in Coho: Culture, Community, Catch), a project supported in part by University Laval, Ressources Aquatiques Québec (RAQ), and the Government of Canada through Genome Canada, Genome British Columbia, and Genome Québec. The funders had no role in study design, data collection and analysis, decision to publish, or preparation of the manuscript.

## Supporting Information Legend

**Table S1: sampling strategy, including various outgroup species.**

**Table S2: Summary Statistics across coho populations, including geographic coordinates, tree branch length, genetic diversity and load statistics**. Figure 1 to Figure 4 can be reconstructed from these.

**Table S3: Outgroup summary statistics by species, with a focus on the load**

**Table S4: Results of linear models.** These provide tests of the relationship between branch length of the population phylogenies based on (a) pairwise Fst value and b) treemix tree inferred with 3 migrations event. c) treemix results obtained after removing the “salmon” river with aberrant branch length

**Table S5: Results of Mann-Whitney test for difference in load among population based on missense mutation and Loss of Function mutation. Mutation effect identified with SNPeff.**

**Table S06: Results of Tukey-HSD test for difference in recombination rate among populations based on Ldhat estimates.** “Pair” is the compared population pair with the initial described in Table S01. “Diff” is the difference in recombination between the pair Lower and Upper CI are the 95% confidence level

**Table S7:** difference in load among diploid region, compared to region of residual tetraploidy with load computed in 500kb windows for each kind of regions

**Tabble S8: AIC of each model for each pairwise comparison.** AM = Ancient Migration, IM = Isolation With Migration, SI = Strict Isolation, SC = Secondary Contact, G = prefix indicating Growth in the descending population, 2N= prefix indicating linked selection, 2m = prefix indicating reduced effective migration along the genome, the A prefix after each model (e.g. AM<u>A</u>) indicated a Growth in the ancestral population

**Table S9: parameter estimates for each of the major regional group where ∂a∂i was fitted.** Na = effective size of the ancestral population, Ne pop1 = effective size of the first population, Nepop2 =effective size of the second population, m12 = migration rate from 2 into 1, m21 being the reverse, me12 = effective migration rate in barriers regions, me21 being the same in the reverse, Tsplit = Split time, Tam = time of migration stop, P = proportion of the genome being neutrally exchanged, Q = proportion the genome undergoing linked selection, O = proportion of the genome correctly oriented, hrf = Hill-Roberston factor, indicating the extent of reduction in Ne in region affected by linked selection, b1, b2 = extend of population growth in current population1 and 2 respectively

**Supplementary Note S1: Reconstruction of demographic history from RADseq data**

**Supplementary Note S2: Relationship between GC3 and recombination**

**Supplementary Note S3 Comparison of Provean and non-synonymous results.**

**Figure S1: PCA and admixture plots among population.**

A Result of a principal component analysis obtained from a set of high quality biallelic SNPs without missing data showing both a clusterization along latitude and longitude as well as discrete clusters corresponding broadly to each river. Each label represents a given individual from a given river (labelled following table S01) and is coloured according to its region of sampling.

B LEA results for admixture inference for K = 14. Each bar represents an individual and is coloured according to its membership probability. Each name corresponds to a river. The TsooYes appear as a mixture of different individuals. Results must be interpreted with caution given the small sample sizes.

**Figure S2: Summary statistics revealed the demography of Coho salmon**

A Positive correlation between the *β*_ST_ and distance to the southernmost site showing that differentiation increased linearly from the south to the north. In all panels each point represents a sampling site and is coloured according to the region in which it was sampled. The most negative values display likely ancestral samples. The Thompson sample displays high inbreeding and is bottlenecked. Displayed is the adjusted R^2^ of a linear model along with its *p-value*. The grey area represents the 95% confidence interval levels around the regression lines obtained with the predict function in R.

B Negative correlation between genetic diversity (observed heterozygosity) and distance to the south.The Thompson sample displays high inbreeding and is bottlenecked. Displayed is the adjusted R^2^ of a linear model along with its *p-value*. The grey area represents the 95% confidence interval levels around the regression lines obtained with the predict function in R.

C Rates of LD decay as a function of distance along the genome. The higher LD indicates a history of inbreeding or bottleneck.

D SMC++ inference of population size change with whole genome sequences for each local population of Coho salmon. Recent times should be interpreted carefully.

**Figure S3: Inference of population split and mixture from Treemix.** A) Tree with three migration arrows. Each name describes a river sample site. Each river is color coded following the color scheme provided elsewhere (e.g. Fig 1). 3 significant migration arrows are displayed. Each migration arrow is colored according to the weight it received (from yellow to red) in Treemix. The weights are related to the fraction of alleles in the descendant population that originated in each donor population. Each node was highly supported based on 500 bootstrap. B) proportion of variance in the covariance of allele frequency explained as a function of the number of migration edges. C) Same tree colored according to the values of πN/πS

**Figure S4: SMC++ split time.** Estimates of population split time from SMC++ under a model without gene flow among populations. Shown are estimates obtained when comparing split time between pairs of samples from different major regional groups. Two different mutation rates were used the: mean and median values based on *Salmo salar* orthologues mapped on the pike *Esox lucius* genome (Wang J. personal communication).

**Figure S5: Correlation between π_N_/π_S_ and π_S_ and π_N_/π_S_ and *N_e_* from smc++** A) Distribution of π_N_/π_S_ as a function of π_S_ in each coho salmon populations from the study. B) Distribution of π_N_/π_S_ as a function of *N_e_*from SMC++ for each coho salmon populations from the study. Results of linear models are displayed. In all panels each point represents a sampling site and is coloured according to the region in which it was sampled. Displayed is the adjusted R^2^ of a linear model along with its *p-value*. The grey area represents the 95% confidence interval levels around the regression lines obtained with the predict function in R.+

**Figure S6:** Correlation between distance to the ocean of each sample location (i.e. corresponding to the spawning migration) and the inferred rate of **A**) non-adaptive substitution (ωNA) and **B**) adaptive substitution (ωA). In all panels each point represents a sampling site and is coloured according to the region in which it was sampled. Displayed is the adjusted R^2^ of a linear model along with its p-value. The grey area represents the 95% confidence interval levels around the regression lines obtained with the predict function in R.

**Figure S7: Results of linear models testing the effect of tree branch length to the root extracted from a *F*_ST_-based population phylogeny on different metrics of selection efficacy A:** relationship between ω_NA_ and tree branch length; **B:** relationship between ω_A_ and tree branch length; **C:** relationship between α and tree branch length. See text for a definition of each metrics. Sample sites are coloured by region. The blue line represents the value of the regression line. In all panels each point represents a sampling site and is coloured according to the region in which it was sampled. Displayed is the adjusted R^2^ along with its p-value.

**Figure S8: Results of linear models testing the effect of tree branch length to the root extracted from a treemix population phylogeny on the load (π_N_/π_S,_ panel A) and different metrics of selection efficacy B:** relationship between ω_NA_ and tree branch length; **C:** relationship between ω_A_ and tree branch length; **D:** relationship between α and tree branch length. See text for a definition of each metrics). Sample sites are coloured by region. The blue line represents the value of the regression line. In all panels each point represents a sampling site and is coloured according to the region in which it was sampled. Displayed is the adjusted R^2^ along with its p-value.

**Figure S9: Correlation between the proportion of amino-acid substitution that results from positive selection (α) and the synonymous diversity π_S_ used as a proxy of effective population size.** Each point represents a sampling site and is coloured according to the region in which it was sampled. Displayed is the adjusted R^2^ along with its p-value. The grey area represents the 95% confidence interval levels around the regression lines obtained with the predict function in R.

**Figure S10: Relationship between GC3 and pN/pS for all populations and all outgroups**

A) Correlation for each population of coho salmon. Each point represents a sampling site and is coloured according to the region in which it was sampled; **B**) correlation within populations of rainbow trout. Each point represent a population as infered using a PCA and corresponds to different rivers of sampling. **C**) correlation for each population of Sockeye and Kokanee ecotype. Each point corresponds to differents rivers.

All correlations are significant. The x-axis displays the median GC3 and y-axis the π_N_/π_S_ ratio. Abbreviation for each site is available in Table S01. Displayed is the adjusted R^2^ of a linear model along with its *p-value*. The grey area represents the 95% confidence interval levels around the regression lines obtained with the predict function in R.

**Figure S11: Relationship between recombination (GC3) and demographic factors (distance to the southernmost site).**

**A)** relationship between the slope of the linear model between GC3 ∼ and π_N_/π_S_ and the distance to the southernmost site. **B)** Correlation between the lowest recombining GC3 classes (expected to display the highest load) and the distance to the southernmost sites. In all panels each point represents a sampling site and is coloured according to the region in which it was sampled. Displayed is the adjusted R^2^ of a linear model along with its *p-value.* The grey area represents the 95% confidence interval levels around the regression lines obtained with the predict function in R.

**Figure S12: region of residual tetraploidy vs diploid region of the genome display different recombination landscapes.** Combined Violin plot and boxplot showing the distribution of population scale recombination (⍴ = 4\**N_e_*\*r) inferred from LDhat in Diploid chromosomes (orange) *vs* the 8 Regions of residual tetraploidy (gray).

**Figure S13: Distribution of recombination rate among all populations**. Each point represents the observed value of population scale recombination rate (ρ = 4*N_e_*\*µ) computed in 1 mb windows over the whole genome in each population. The harmonic mean is plotted by a red dot along with its value. For each population a violin plot embedded within a boxplot is shown.

**Figure S14: TE length differs between diploid chromosome versus chromosome displaying residual tetraploidy.** Violin plot displaying the difference in TEs relative length (i.e. length corrected by the total length of each chromosome) for each major TE category and each type of chromosome. Red point = mean +/− 1*sd.

**Figure S15: Provean analysis of potentially deleterious mutation and the resulting load Boxplot showing the number of deleterious alleles per river (sorted from the south to to north)**

*left panel = additive load, right panel= recessive load*

*top = missense deleterious mutations according to Provean predictions, bottom = missense tolerated mutations according to Provean predictions*

No strong differences are observed in the additive load among populations.

Significant differences were observed for the recessive load in populations at the expansion front which is qualitatively similar to our inferences from missense and LoF mutations. Each color represents a major regional group. Results were obtained for a random subset of mutations only given the strong computational burden of Provean.

