## Supplemental material for "Allele surfing causes maladaptation in a Pacific salmon of conservation concern"

#### Supplementary Note S1: Reconstruction of demographic history from RADseq data

Previous work [1] based on ∂a∂i [2] and Fastsimcoal [3] suggested the existence of two historical refugia one located in the South of the distribution range, another that would correspond to a refugia for the Thompson lineage which is strongly differentiated, based on RADseq and whole genome sequences. Inferences also suggested that the Thompson area would have diverged approximately ~100 to 130 kyA followed and became reconnected in postglacial time. Yet, levels of net sequence divergence and of genetic diversity as well as the presence of long runs of homozygosity (ROH) are compatible with a recent bottleneck and more recent divergence of this population from the rest of the dataset[1,4].

Moreover, SMC++ analysis in [4], suggests a more recent divergence (in the range 5-60kyA). Therefore we aimed at validating the results from our precedent work suggesting that *i*) populations have expanded from a single refugia in postglacial time and *ii)* the Thompson have been isolated in post glacial time and then undergone a secondary contact [1].

The major novelty of our modelling approach is to include the possibility for population size change (expansion, bottleneck) in the ancestral population, in addition to size change in the daughter populations. Indeed a recent simulation study suggested that secondary contact may falsely be inferred under a true model of continuous migration if ancestral population size change is ignored [5]. In addition, we took advantage of our outgroup information to build unfolded site frequency spectra, providing richer information than our previous inferences [1]. As previously, our models included the effect of linked selection and barriers to gene flow which are respectively modeled through variation in effective population size and variation in effective migration rate. These models were applied to the exact same set of RADseq data [1], but based on a new unfolded joint site frequency spectrum constructed after remapping all data to the V2 genome. All unfolded jSFS were generated with ANGSD [6]. Model choice was performed using AICc.

We fitted four alternative models of divergence, described in [1] using ∂a∂i [2]:

1) Strict Isolation (SI)

2) Divergence with Ancient Migration (AM)

3) Isolation with Migration (IM)

4) Divergence followed by Secondary Contact (SC).

These models integrate the confounding effect of linked selection modeled as a reduction in effective population along the genome (suffix 2N) as well as barriers to gene flow reducing the local migration rate along the genome (suffix 2M).

As previously we compared the following major genetic group by fitting pairwise models of populations divergence for 5 pairs of populations:

* California *vs* Thompson

* Cascadia *vs* Thompson

* Alaska *vs* Thompson

* California *vs* Alaska

* Cascadia *vs* Alaska

#### Results

Model choice performed through AIC and ΔAIC revealed support for the model of ancient migration (AM) in all pairwise comparisons (**Table S08**, **Table S09**). In 3 out of the 5 comparisons, models that integrated population size change in the ancestral population received higher support. Models with either linked selection (suffix 2N) or barriers to gene flow (suffix 2M) were always supported compared to models that ignore these effects. Importantly, these models provided a better fit to the data than our previous work in terms of residual as shown in the figure below. Suggesting that our new inferences should be more reliable both in terms of model choice and parameter estimation. Parameter estimates are provided in **Table S09**. Our main parameter of interest here, the split time revealed that the populations have diverged between 5KyA and 45 KyA, which is more recent than our previous work. This result is more in line with SMC++ based inference (**Fig S03**) despite the very different modelling assumption of the two approaches.

In summary, our results do not support the existence of a separate refuge followed by secondary contact, but rather support a recent divergence with initial gene flow.

***Figure Supp Note S1:*** **Model and Residual for each pairwise comparison obtained from ∂a∂i.** Each plot displays the observed jSFS (data), the modeled jSFS (model) and the residuals. The best model was inferred using ΔAIC and AIC weights (Table S2).

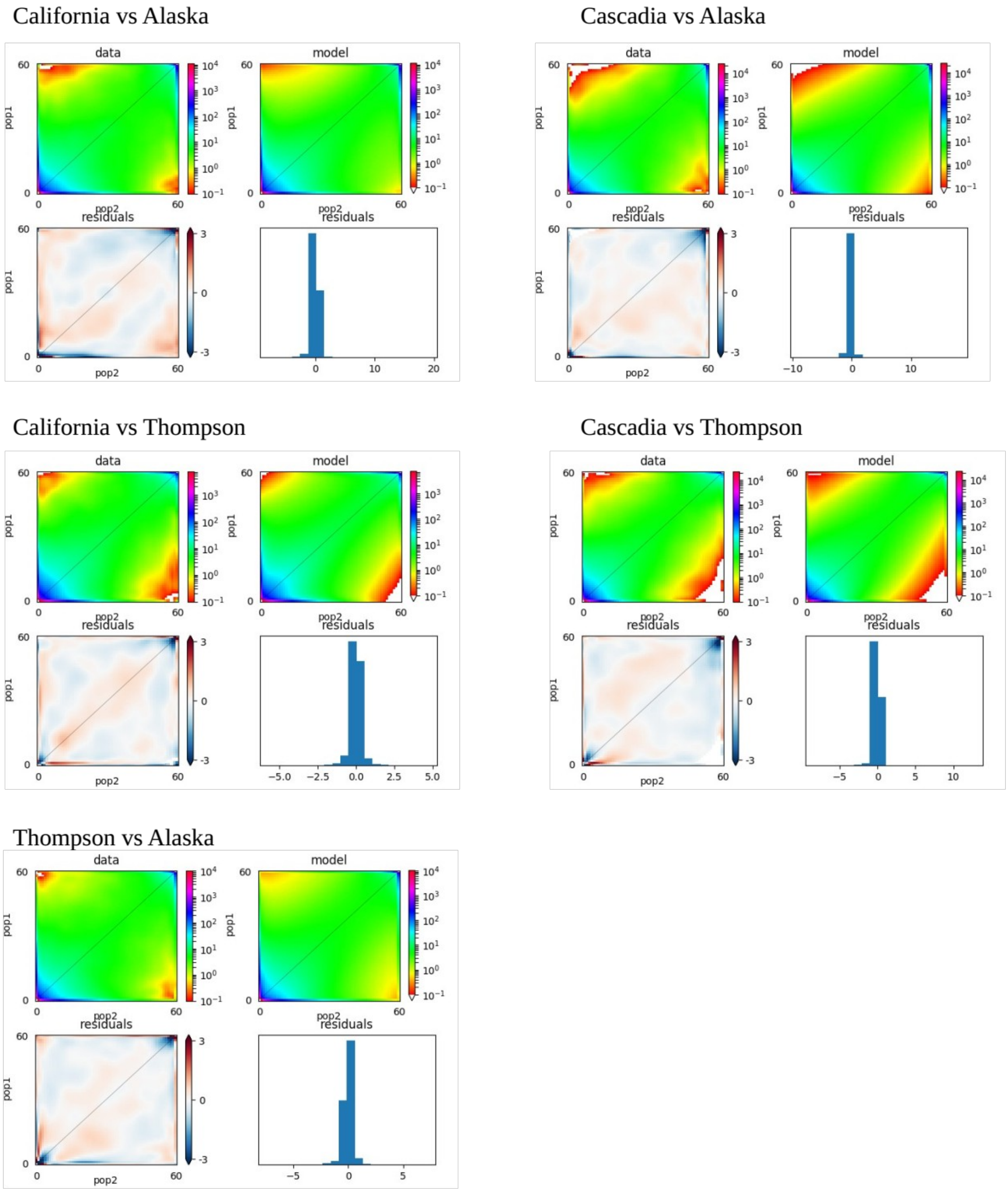

#### Supplementary Note S2: Relationship between GC3 and recombination

Instead of directly testing the association between $\pi_N/\pi_S$ and recombination, we assessed the relationship between $\pi_N/\pi_S$ and GC content at the third coding position (GC3). There were four reasons for that. First, population genetic estimates relies on the compound metric $4N_e$*rho, rather than rho directly; so that strong variation in effective population size is a first source of bias. Second, gene flow is known to distort these results [7]. Third, under some conditions of mutation rates or low recombination, these methods may be biased [8]. Fourth, Ldhat and several related methods often assumed demographic equilibrium, a hypothesis that was strongly violated at the expansion front or in the Salmon river, showing signs of bottleneck and inbreeding.

Indeed, during the recombination process, GC biased gene conversion occurs (gBGC): during recombination, heteroduplexes mispairing are preferentially repaired in favor of G-C based, rather than A-T, this is the process of gGBC. The higher the recombination is, the higher the probability of errors is, and therefore an increase in GC content is to be expected, as observed in several species [9–16].

Unlike population scale estimates of recombination, GC3 estimates do not rely on a particular population genetic model that assumes demographic equilibrium, it should therefore not be biased by the process mentioned above (gene flow, variation in population size). It can be directly estimated from the data and the accuracy of measuring GC3 should not be particularly affected by variation in mutation or recombination rates. Finally, the GC3 should remain a good proxy although ideally, we should use GC3 at silent sites (GC3s) that are even less affected by selection [13], or 4-fold degenerated GC3.

#### Supplementary Note S3 Comparison of Provean and non-synonymous results.

Provean [17] measures the effect of a variation (*v*) on a protein A by measuring the similarity (using blast) of the protein A to a protein B before and after the introduction of the variation. The assumption is that a variation that reduces the similarity of protein A to the functional homolog protein B is more likely to cause a damaging effect. To this end, a change in the “alignment score” (delta score) can be used as a measure of change in “similarity” caused by a variation. A low delta score is interpreted as a deleterious effect of the variation, with a score below -2.5 being the recommended value to provide a tradeoff between specificity (i.e. a high confidence in the variants being identified as deleterious) and sensitivity (i.e. identifying a high number of deleterious variants). A high delta score is interpreted as neutral. The top 30 clusters of closely related proteins are used to compute a delta score, which are then averaged across clusters [17]

While we previously used Provean to identify potentially deleterious mutations, we argue here that such an approach, based on the protein conservation across many species, is similar in spirit to approaches such as GERPP score [18], whose population genetic properties have been well studied [19]. In particular, a major limit of these approaches is associated with their assumption of conservation across all the clades investigated, so that only mutation that was selected across all the studied clades will produce a significant score. Change in selection at a given site through time reduces the power to detect selection in our focal species. Moreover, these approaches were mostly designed for humans and mammals. Still, to verify the consistency of our results we run Provean on a random set of non-synonymous gene. The results provided in **Figure S13** remained broadly consistent with those obtained from LoF and missense mutations in Figure 3.

### Supplementary Figures

**Figure S1: PCA and admixture plots among population.**

- **A** Result of a principal component analysis obtained from a set of high quality biallelic SNPs without missing data showing both a clusterization along latitude and longitude as well as discrete clusters corresponding broadly to each river. Each label represents a given individual from a given river (labelled following table S01) and is coloured according to its region of sampling.
- B LEA results for admixture inference for K = 14. Each bar represents an individual and is coloured according to its membership probability. Each name corresponds to a river. The TsooYes appear as a mixture of different individuals. Results must be interpreted with caution given the small sample sizes.

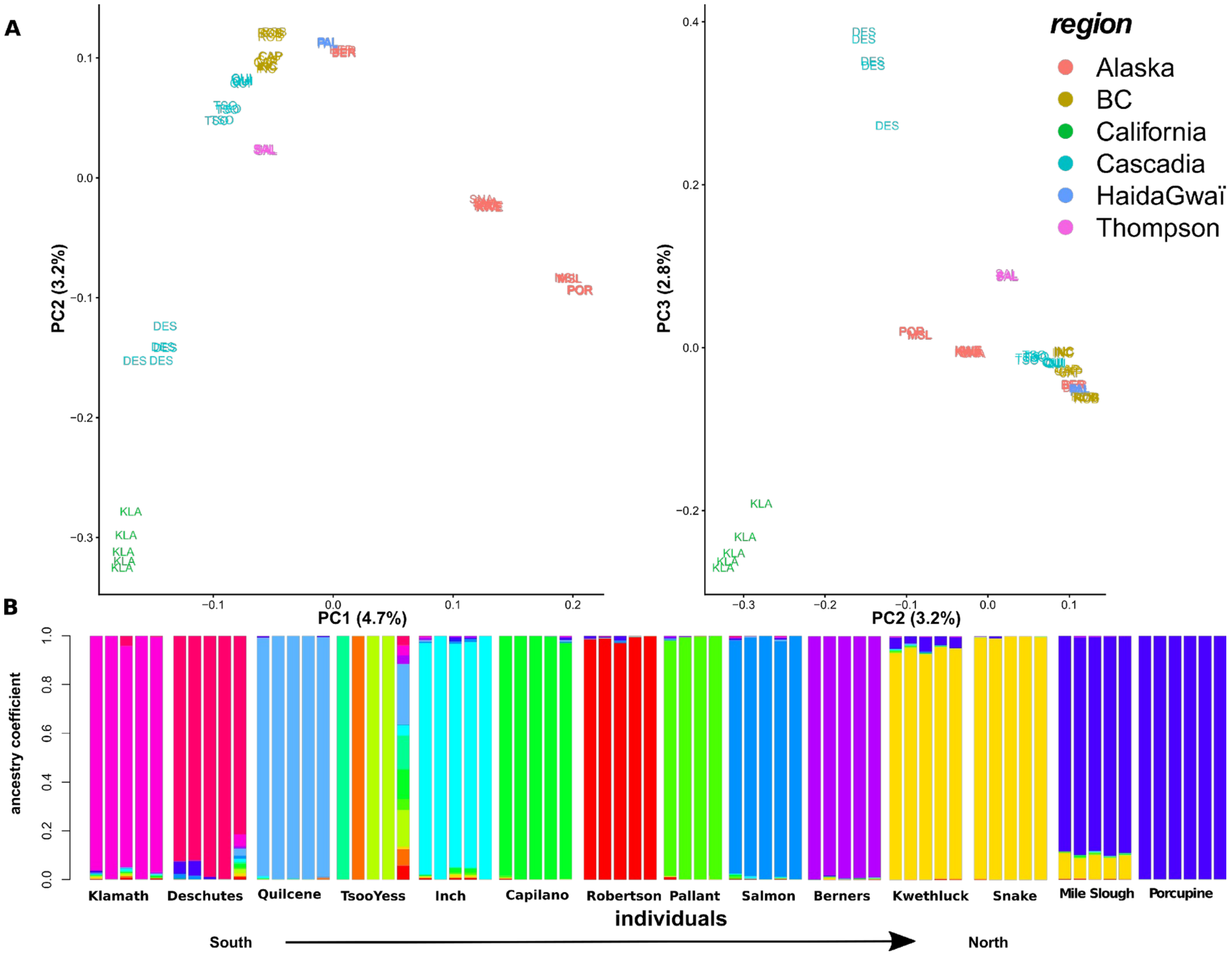

**Figure S2: Summary statistics revealed the demography of Coho salmon**

A Positive correlation between the $\beta_{ST}$ and distance to the southernmost site showing that differentiation increased linearly from the south to the north. In all panels each point represents a sampling site and is coloured according to the region in which it was sampled. The most negative values display likely ancestral samples. The Thompson sample displays high inbreeding and is bottlenecked [1,4]. Displayed is the adjusted $R^2$ of a linear model along with its *p-value*. The grey area represents the 95% confidence interval levels around the regression lines obtained with the predict function in R.

B Negative correlation between genetic diversity (observed heterozygosity) and distance to the south.The Thompson sample displays high inbreeding and is bottlenecked. Displayed is the adjusted $R^2$ of a linear model along with its *p-value*. The grey area represents the 95% confidence interval levels around the regression lines obtained with the predict function in R.

C Rates of LD decay as a function of distance along the genome. The higher LD indicates a history of inbreeding or bottleneck.

D SMC++ inference of population size change with whole genome sequences for each local population of Coho salmon. Recent times should be interpreted carefully.

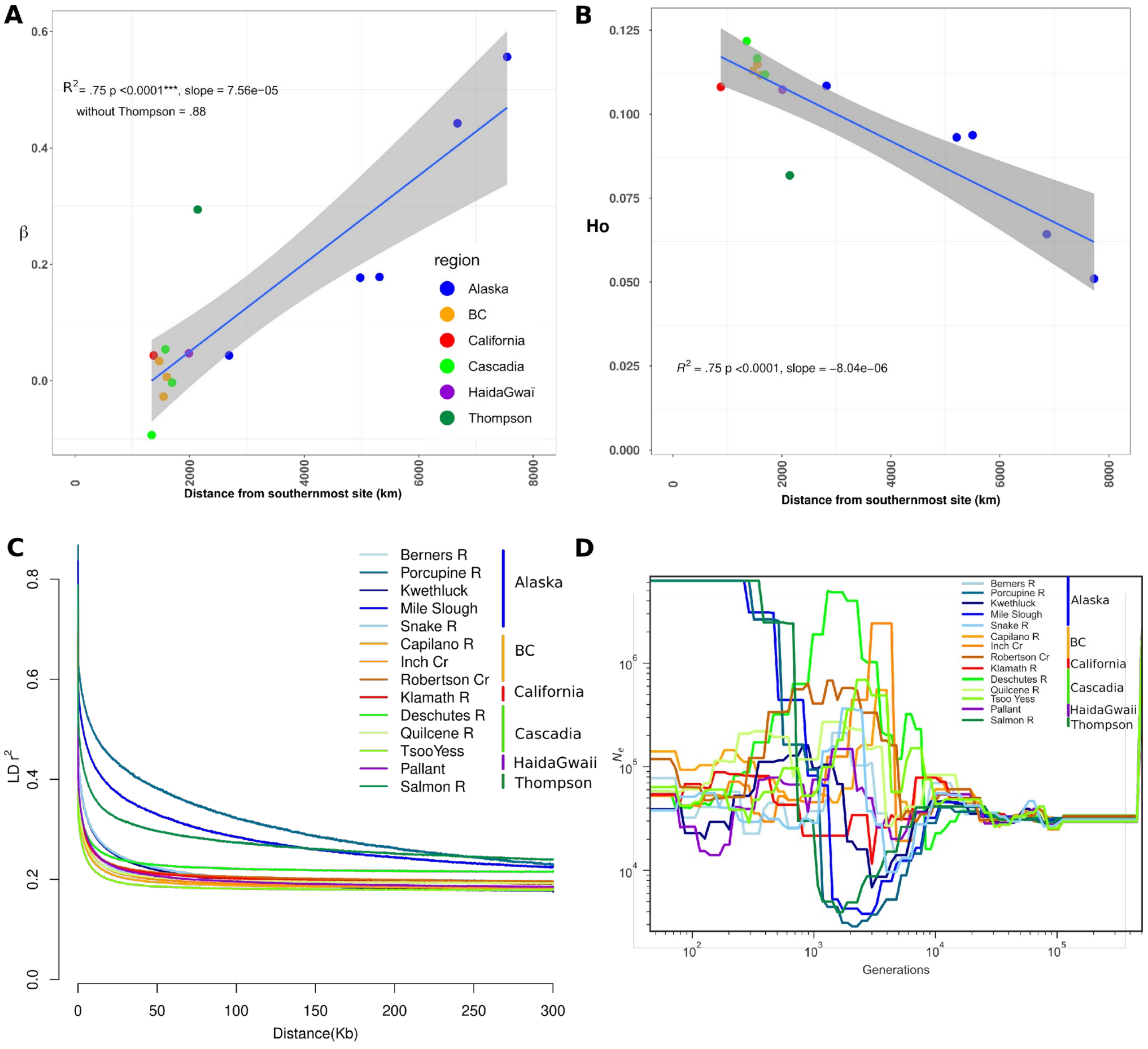

**Figure S3: Inference of population split and mixture from Treemix.**

**A)** Tree with three migration arrows. Each name describes a river sample site. Each river is color coded following the color scheme provided elsewhere (e.g. Fig 1)**.** 3 significant migration arrows are displayed. Each migration arrow is colored according to the weight it received (from yellow to red) in Treemix. The weights are related to the fraction of alleles in the descendant population that originated in each donor population. Each node was highly supported based on 500 bootstraps. **B)** proportion of variance in the covariance of allele frequencies explained as a function of the number of migration edges. **C)** Same tree colored according to the values of $\pi_N/\pi_S$

**A** **B**

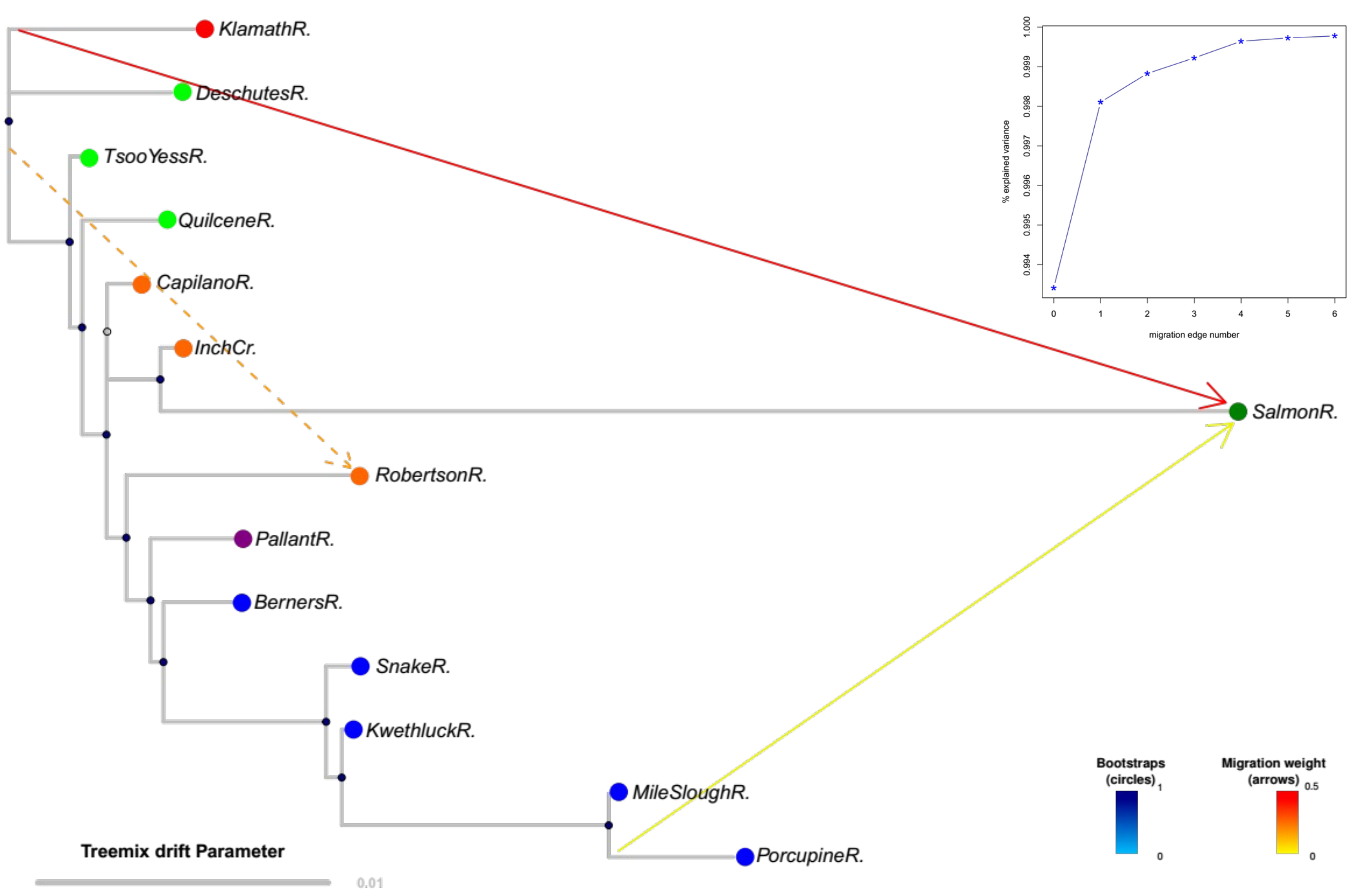


**C**

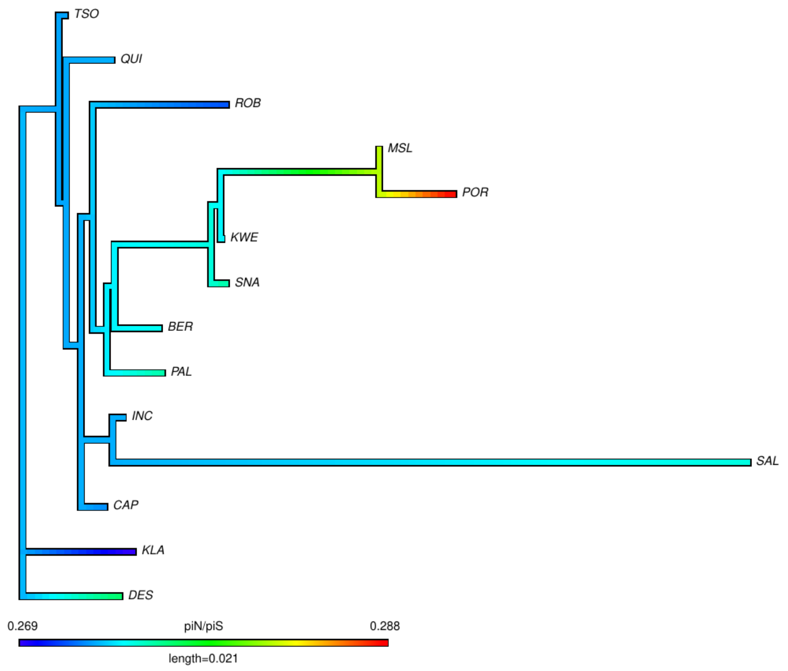

**Figure S4: SMC++ split time.**

Estimates of population split time from SMC++ under a model without gene flow among populations. Shown are estimates obtained when comparing split time between pairs of samples from different major regional groups. Two different mutation rates were used: the mean and median values based on *Salmo salar* orthologues mapped on the pike *Esox lucius* genome (Wang J. personal communication).

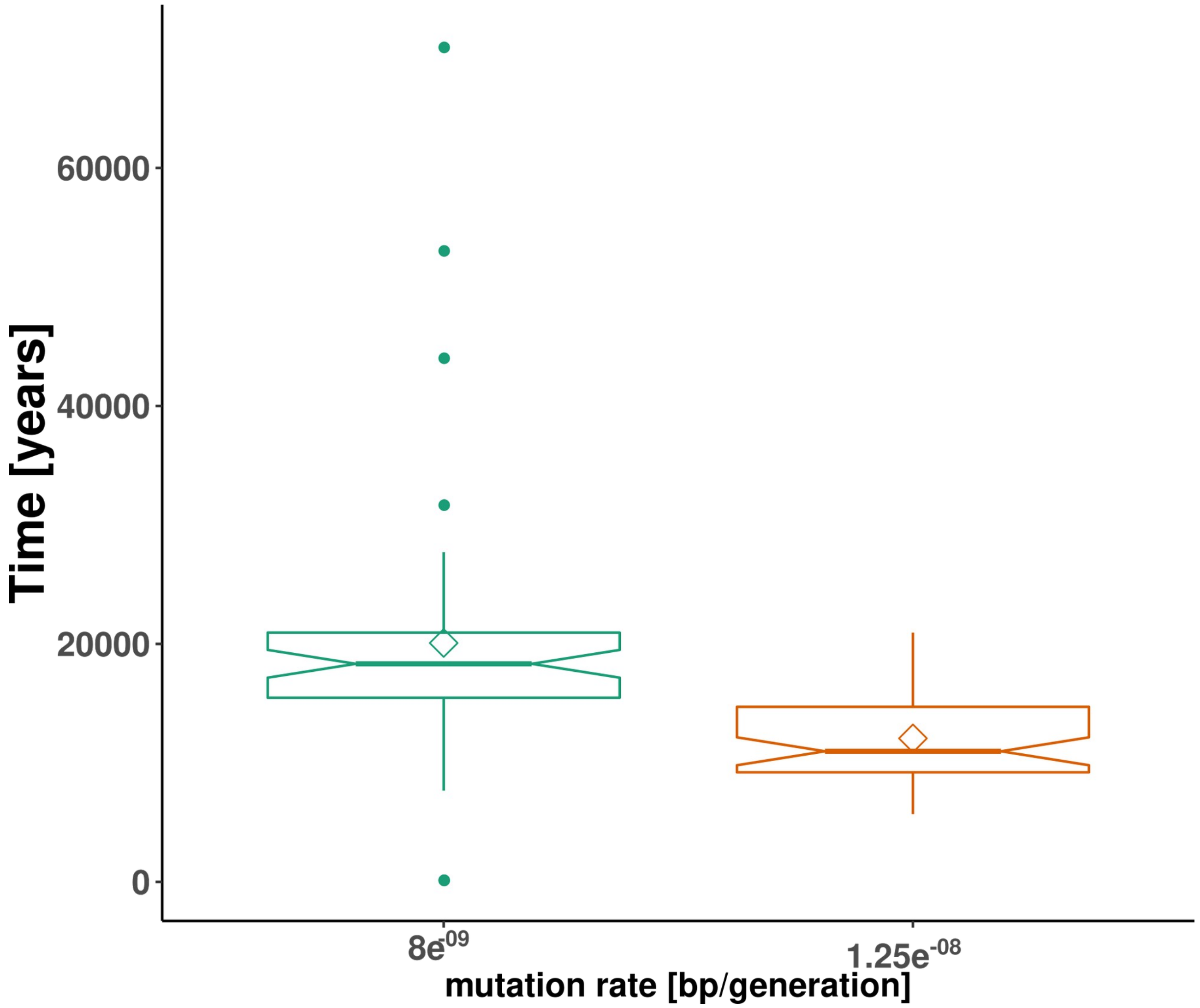

**Figure S5: Correlation between $\pi_N/\pi_S$ and $\pi_S$ and $\pi_N/\pi_S$ and $N_e$ from smc++**

**A**) Distribution of $\pi_N/\pi_S$ as a function of $\pi_S$ in each Coho salmon populations from the study. **B**) Distribution of $\pi_N/\pi_S$ as a function of $N_e$ from SMC++ for each Coho salmon populations from the study. Results of linear models are displayed. In all panels each point represents a sampling site and is coloured according to the region in which it was sampled. Displayed is the adjusted $R^2$ of a linear model along with its *p-value*. The grey area represents the 95% confidence interval levels around the regression lines obtained with the predict function in R.+

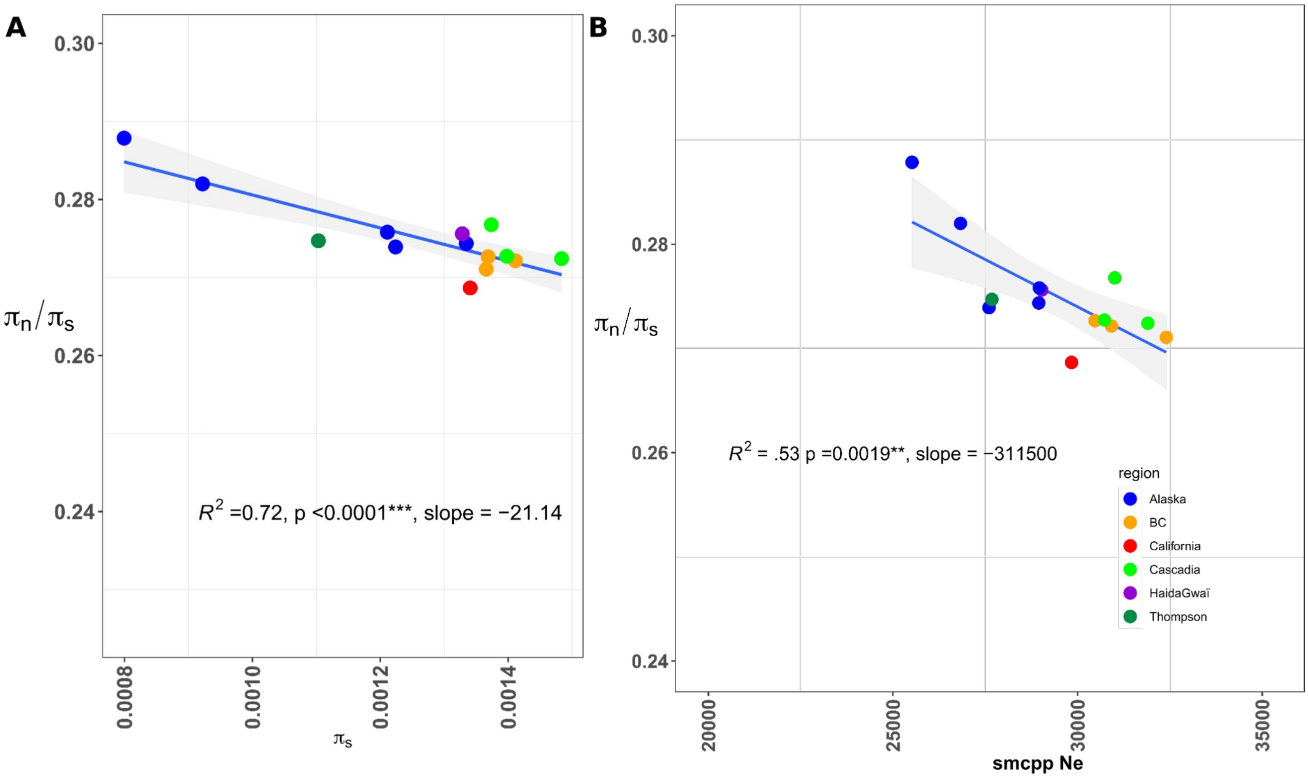

**Figure S6:** Correlation between distance to the ocean of each sample location (i.e. corresponding to the spawning migration) and the inferred rate of **A**) non-adaptive substitution ($\omega_{NA}$) and **B**) adaptive substitution ($\omega_A$). In all panels each point represents a sampling site and is coloured according to the region in which it was sampled. Displayed is the adjusted $R^2$ of a linear model along with its *p-value*. The grey area represents the 95% confidence interval levels around the regression lines obtained with the predict function in R.

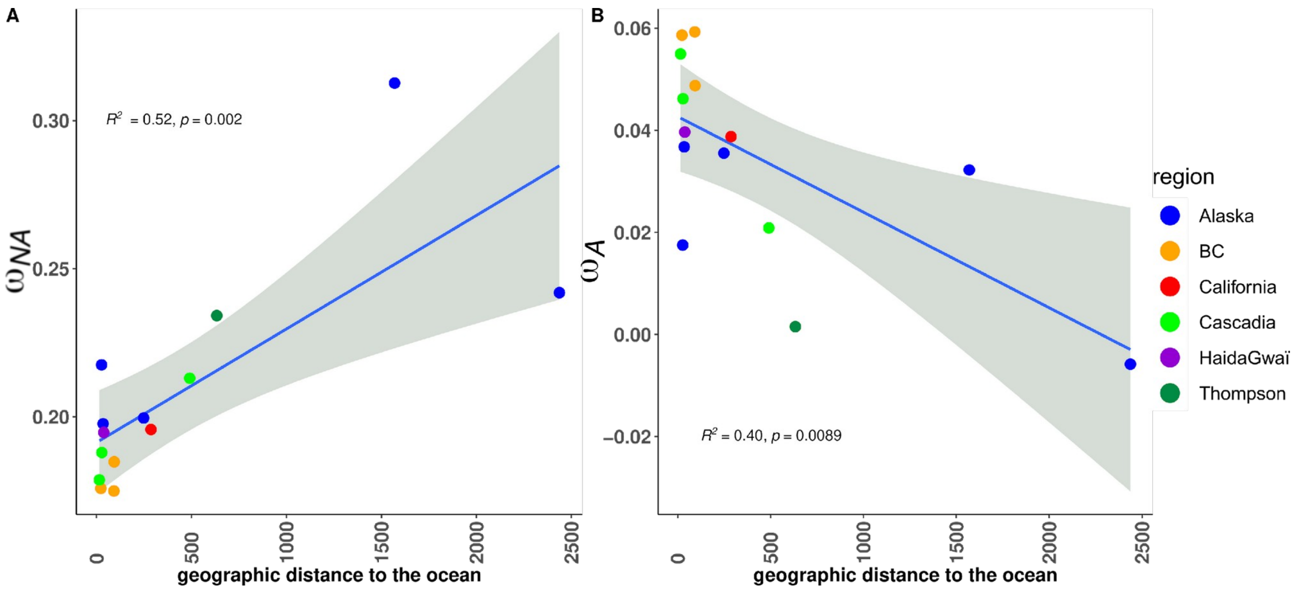

**Figure S7: Results of linear models testing the effect of tree branch length to the root extracted from a $F_{ST}$-based population phylogeny on different metrics of selection efficacy. A:** relationship between $\omega_{NA}$ and tree branch length; **B:** relationship between $\omega_A$ and tree branch length; **C:** relationship between $\alpha$ and tree branch length. See text for a definition of each metrics**.** Sample sites are coloured by region. The blue line represents the value of the regression line. In all panels each point represents a sampling site and is coloured according to the region in which it was sampled. Displayed is the adjusted $R^2$ of a linear model along with its *p-value*.

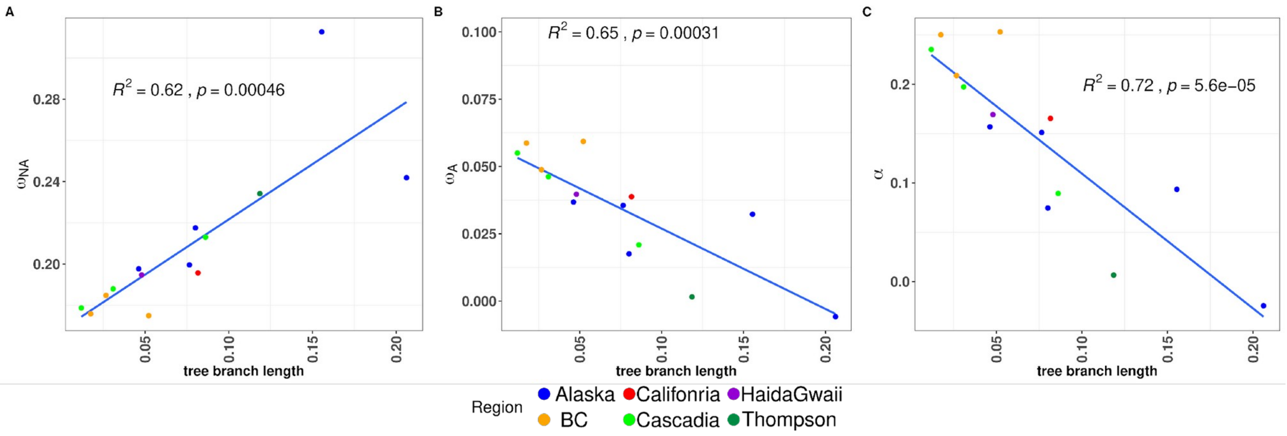

**Figure S8: Results of linear models testing the effect of tree branch length to the root extracted from a treemix population phylogeny on the load ($\pi_N/\pi_S$, panel A) and different metrics of selection efficacy. B:** relationship between $\omega_{NA}$ and tree branch length; **C:** relationship between $\omega_A$ and tree branch length; **D:** relationship between $\alpha$ and tree branch length. See text for a definition of each metrics)**.** Sample sites are coloured by region. The blue line represents the value of the regression line. In all panels each point represents a sampling site and is coloured according to the region in which it was sampled. Displayed is the adjusted $R^2$ of a linear model along with its *p-value*.

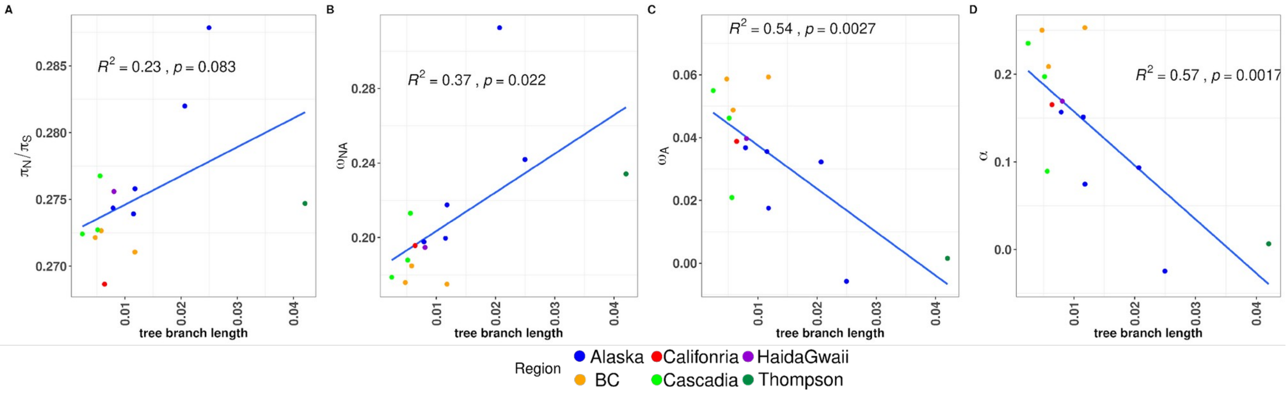

**Figure S9: Correlation between the proportion of amino-acid substitution that results from positive selection (α) and the synonymous diversity $\pi_S$ used as a proxy of effective population size.**

Each point represents a sampling site and is coloured according to the region in which it was sampled. Displayed is the adjusted $R^2$ of a linear model along with its *p-value*

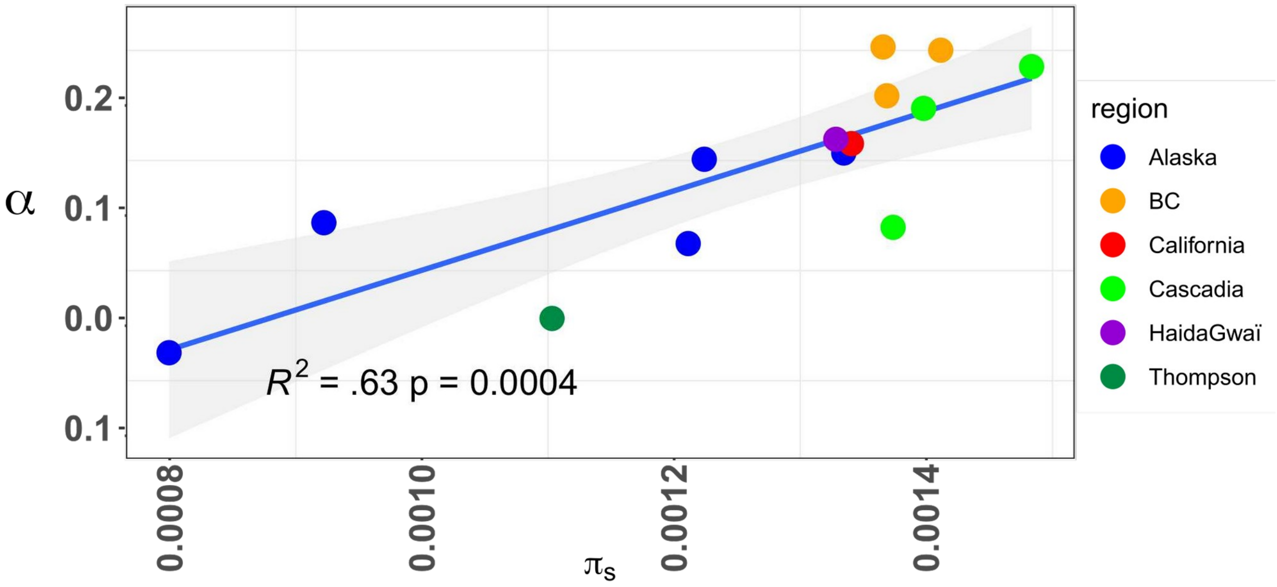

**Figure S10: Relationship between GC3 and pN/pS for all populations and all outgroups**

A) Correlation for each population of coho salmon. Each point represents a sampling site and is coloured according to the region in which it was sampled; **B**) correlation within populations of rainbow trout. Each point represent a population as infered using a PCA and corresponds to different rivers of sampling. **C**) correlation for each population of Sockeye and Kokanee ecotype. Each point corresponds to differents rivers. All correlations are significant. The x-axis displays the median GC3 and y-axis the $\pi_N/\pi_S$ ratio. Abbreviation for each site is available in Table S01. Displayed is the adjusted $R^2$ of a linear model along with its *p-value*. The grey area represents the 95% confidence interval levels around the regression lines obtained with the predict function in R.

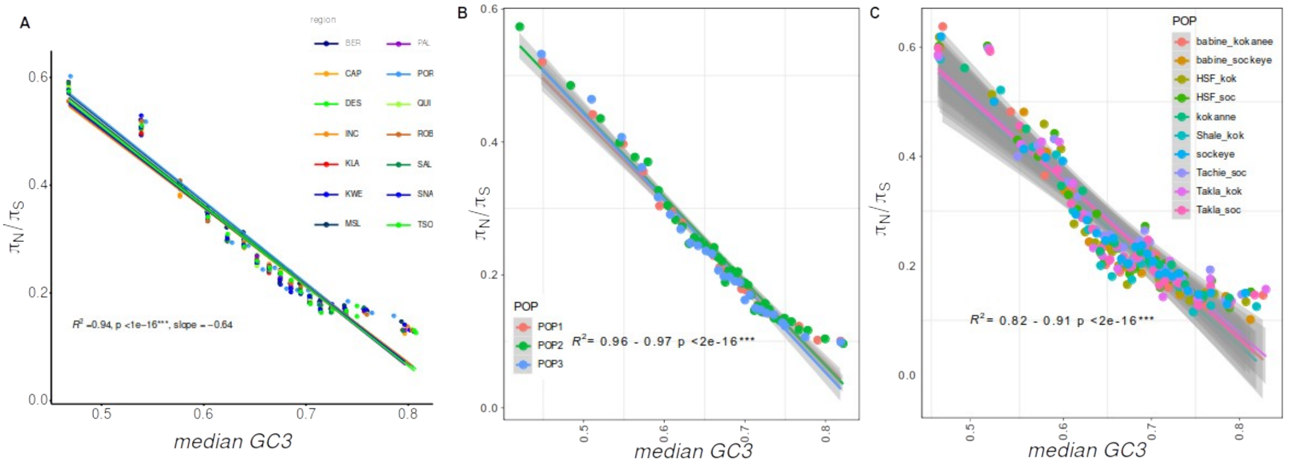

**Figure S11: Relationship between recombination (GC3) and demographic factors (distance to the southernmost site).**

**A)** relationship between the slope of the linear model between GC3 ~ and $\pi_N/\pi_S$ and the distance to the southernmost site. **B)** Correlation between the lowest recombining GC3 classes (expected to display the highest load) and the distance to the southernmost sites. In all panels each point represents a sampling site and is coloured according to the region in which it was sampled. Displayed is the adjusted $R^2$ of a linear model along with its *p-value*. The grey area represents the 95% confidence interval levels around the regression lines obtained with the predict function in R.

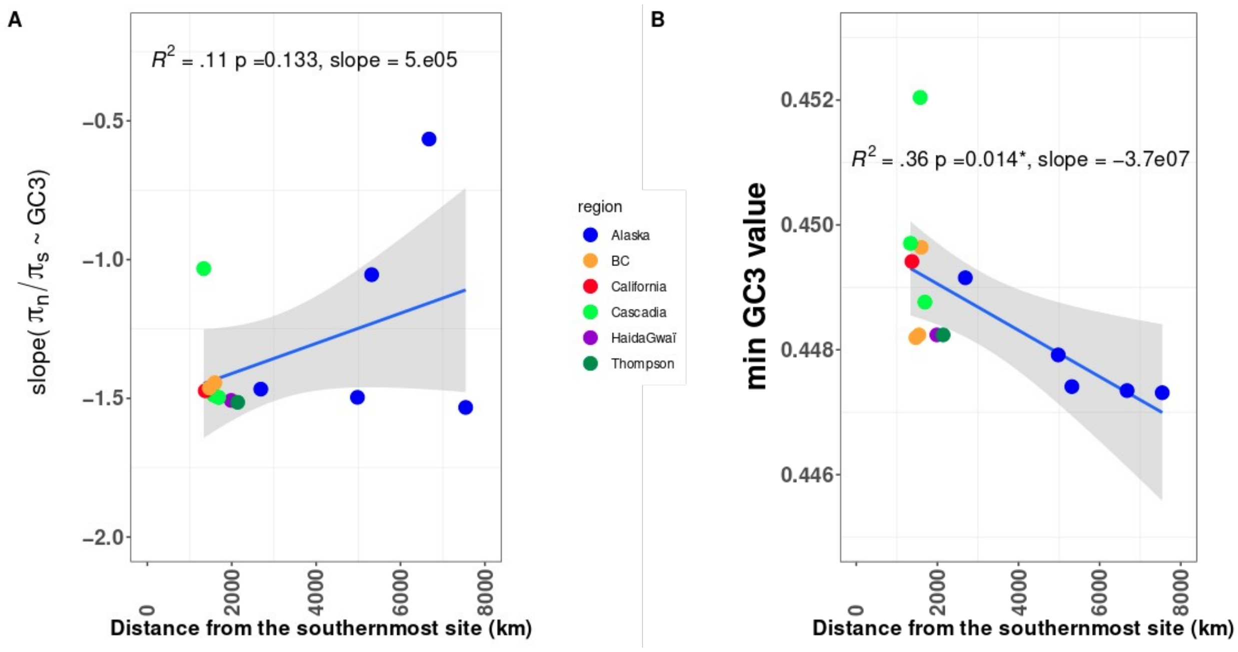

**Figure S12: Regions of residual tetraploidy vs diploid regions of the genome display different recombination landscapes.**

Combined Violin plot and boxplot showing the distribution of population scale recombination ($\rho = 4*N_e*r$) inferred from LDhat in Diploid chromosomes (orange) *vs* the 8 Regions of residual tetraploidy (gray).

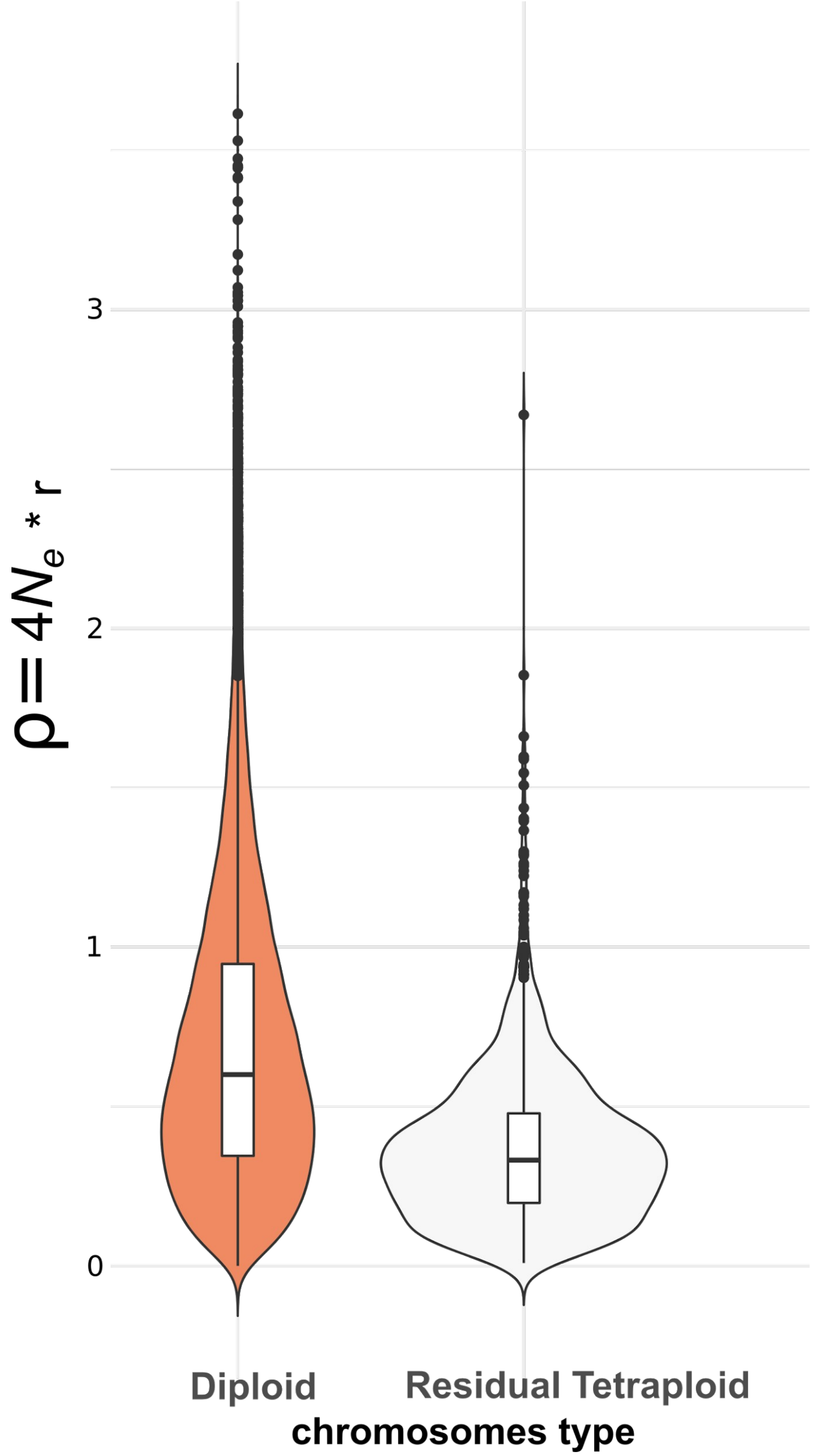

**Figure S13: Distribution of recombination rate among all populations.** Each point represents the observed value of population scale recombination rate (ρ = 4$N_e$*μ) computed in 1 mb windows over the whole genome in each population. The harmonic mean is plotted by a red dot along with its value. For each population a violin plot embedded within a box-plot is shown.

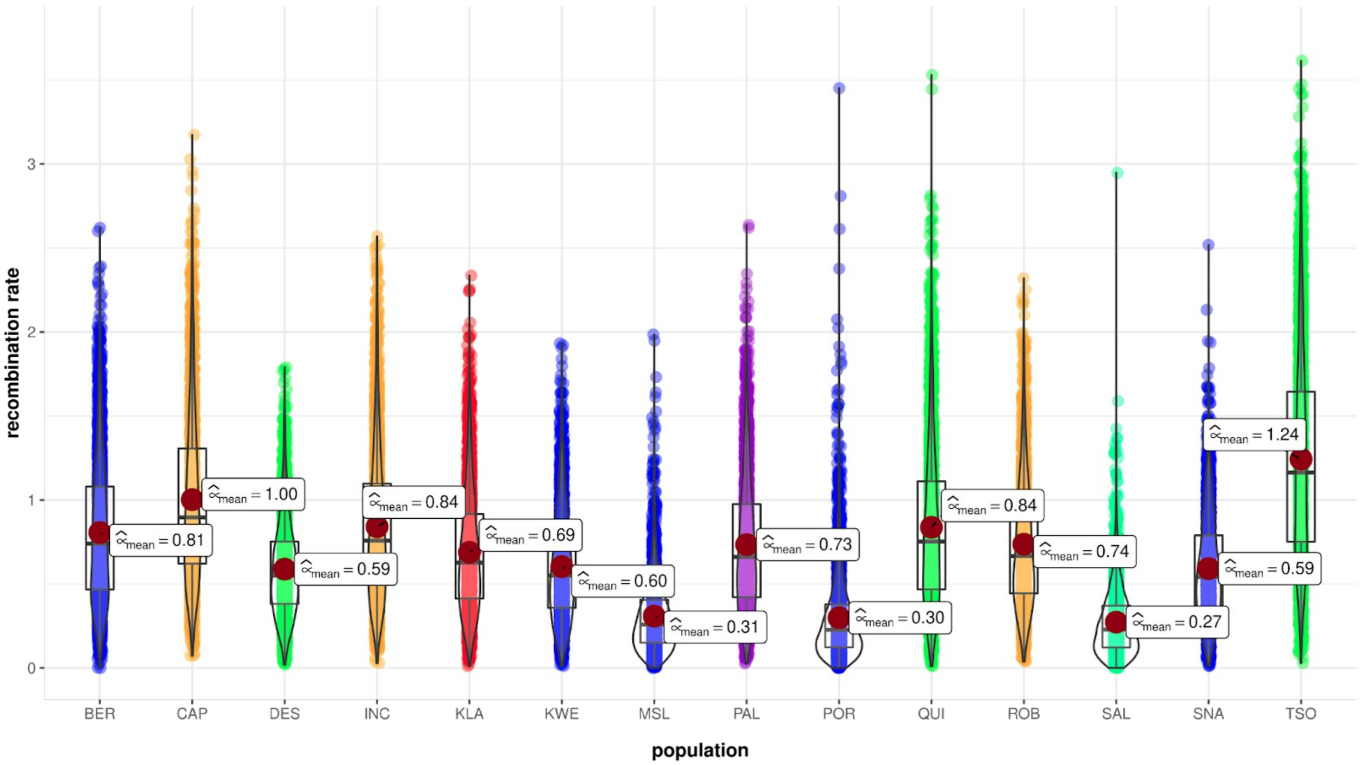

**Figure S14: TE length differs between diploid chromosome versus chromosome displaying residual tetraploidy.**

Violin plot displaying the difference in TEs relative length (i.e. length corrected by the total length of each chromosome) for each major TE category and each type of chromosome. Red point = mean +/- 1*sd.

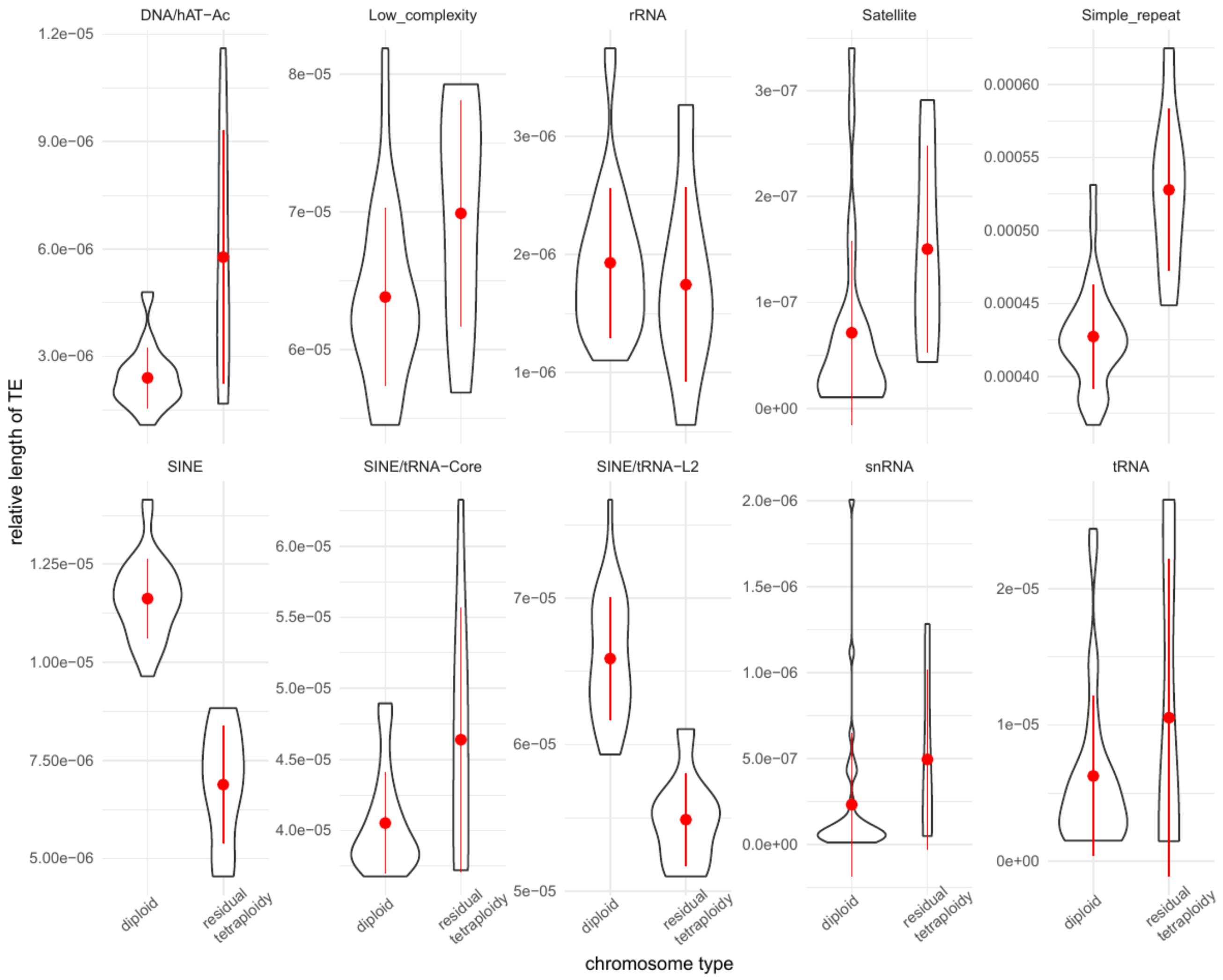

**Figure S15: Provean analysis of potentially deleterious mutation and the resulting load**
***Boxplot showing the number of deleterious alleles per river (sorted from the south to to north)***

*left panel = total load*
*right panel= recessive load*
*top = missense deleterious mutations according to Provean predictions*
*bottom = missense tolerated mutations according to Provean predictions*
No strong differences are observed in the total load among populations.
Significant differences were observed for the fixed load in populations at the expansion front which is qualitatively similar to our inferences from missense and LoF mutations.
Each color represents a major regional group.
Results were obtained for a random subset of mutations only given the strong computational burden of Provean.

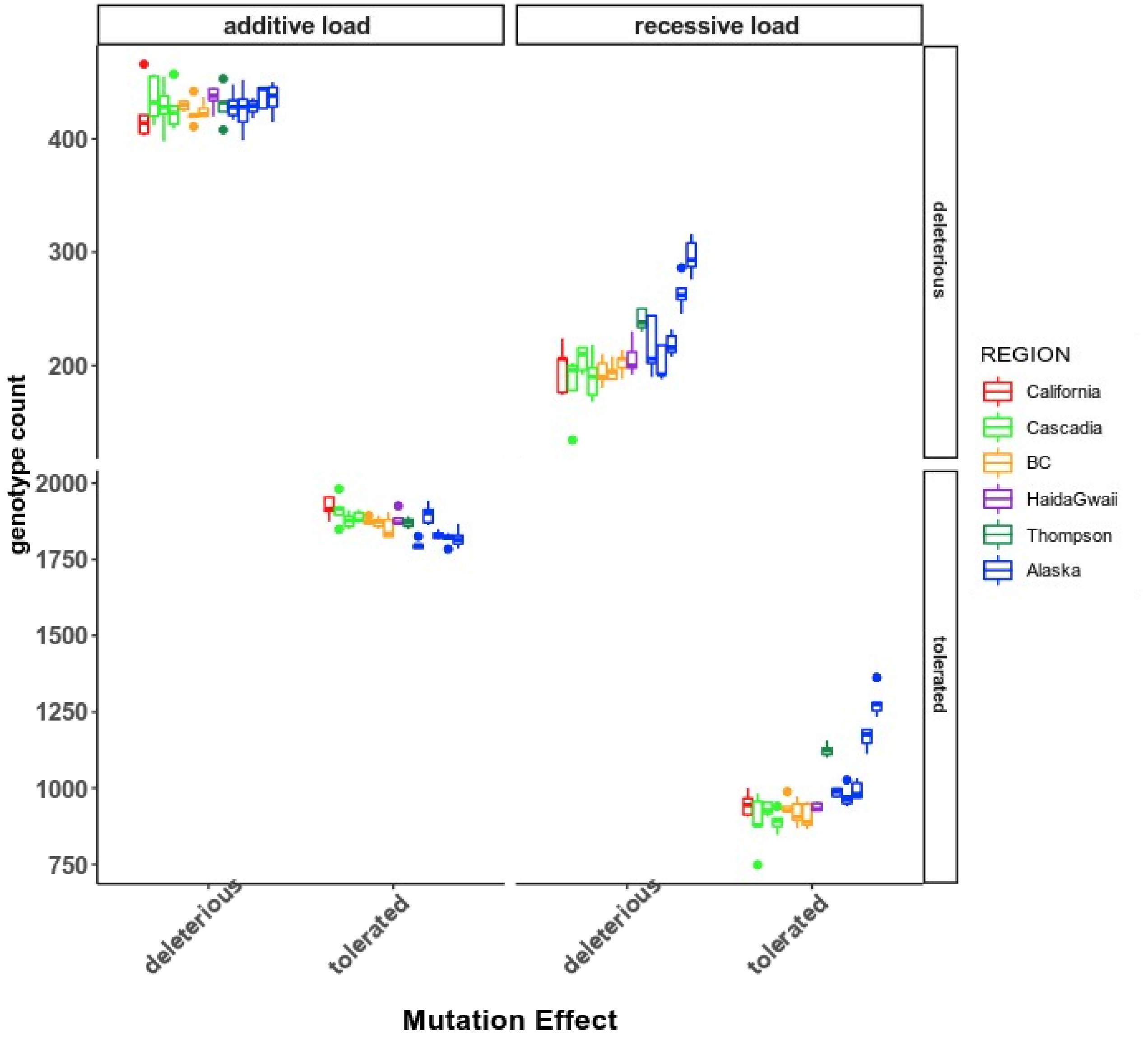

#### Pipeline availability:

Whole genome SNP Calling:
https://github.com/QuentinRougemont/gatk_haplotype

SMC++ inference:
https://github.com/QuentinRougemont/smcpp_input

Recombination inference:
https://github.com/QuentinRougemont/LDhat_workflow,

∂a∂i reconstruction:
https://github.com/QuentinRougemont/DemographicInference;

Deleterious mutation identification and load:
https://github.com/QuentinRougemont/piNpiS
https://github.com/QuentinRougemont/provean

Other analysis (tajima's D, betaST, he, ho, genetic structure, PCA):
https://github.com/QuentinRougemont/utility_scripts
